# Origin of Class B J-domain proteins involved in amyloid transactions

**DOI:** 10.1101/2025.09.06.674649

**Authors:** Przemyslaw Domanski, Milena Stolarska, Katarzyna Kalinowska, Dominik Purzycki, Brenda A. Schilke, Hubert Wyszkowski, Marcin Pitek, Aneta Szymanska, Katarzyna Bury, Jacek Czub, Agnieszka Kłosowska, Elizabeth A. Craig, Jaroslaw Marszalek, Bartlomiej Tomiczek

## Abstract

J-domain protein (JDP) chaperones function widely in proteostasis. Notably, eukaryotic class B JDPs of the cytosol/nucleus prevent assembly or drive disassembly of amyloid aggregates known to cause neurodegenerative diseases, yet their evolutionary origin is not known. Members of the most ubiquitous class B subgroup, canonical B (B^C^) JDPs, lack the signature zinc finger region (ZnF) of the more prevalent class A JDPs, while having other key features in common. Our phylogenetic analysis revealed that B^C^ JDPs evolved more than once from class A duplicates, losing their ZnF. The cytonuclear B^C^s emerged at the base of eukaryotes. Cytonuclear class B’ (i.e. B’^(ST)^) JDPs that have a substrate binding domain of unknown origin, distinct from that of As and B^C^s, emerged from a B^C^ duplication at the base of metazoans and subsequently multiplied by duplications. Origin of B’^(ST)^s, which are capable of suppressing formation of amyloid aggregates, predated the emergence of disease-causing amyloidogenic proteins. Using ancestral sequence resurrection, we tested when cytonuclear Bs evolved their amyloid related functions. We found that their common ancestor with As, AncAB that has a ZnF does not facilitate disassembly of amyloid fibrils, while AncB, which lacks a ZnF, is active. Overall, our findings are consistent with the idea that, though the ZnF of class A JDPs is important for some roles, its loss allowed evolution of novel functions, as illustrated by the ability of B^C^ and B’^(ST)^ JDPs to control amyloid aggregate levels.

**Significance statement:** Across procaryotes and eukaryotes J-domain proteins (JDPs) are key players in Hsp70 chaperone systems that maintain cellular protein homeostasis. The abundant class A and B JDPs have structural similarities, yet their origin has remained unresolved. Here we show that B JDPs independently evolved from As more than once. In each case A lost its zinc finger (ZnF) domain, suggesting that such loss has allowed evolution of new functions. Supporting this idea, biochemical resurrection of an ancestral eukaryotic B revealed that its ability to disassemble amyloid aggregates, differentiating it from As, evolved after ZnF loss. Later, the subset of Bs implicated in suppression of disease-causing amyloid aggregate formation originated from a duplicate of this B in the common ancestor of animals.

## Introduction

J-domain proteins (JDPs) are obligatory cochaperones of Hsp70s. Together, JDP/Hsp70 systems function in many cellular processes – from protein homeostasis, including folding of polypeptide chains and maintenance of protein structure upon stress, to remodelling of protein complexes and aggregates, including amyloids responsible for human disease (1-8). Interaction of Hsp70 with a substrate polypeptide is critical for all these functions (9). All JDPs have a J-domain that interacts with Hsp70, stimulating its inherently low ATPase activity, thereby stabilizing substrate interaction (10, 11). Many, including the first identified and best-studied JDP, DnaJ of *Escherichia coli*, also bind substrates (12, 13). Via the J-domain-Hsp70 interaction, these JDPs “deliver” their bound substrates to their Hsp70 partners (14). Together with a nucleotide exchange factor (NEF), JDPs and Hsp70s form systems for efficient cycles of interaction with substrate protein. By prioritizing substrate interaction in this way, JDPs can be said to “direct” Hsp70s to perform specific function(s) in cells (1, 2, 6, 7).

Hundreds of structurally and functionally divergent JDPs have been identified across the tree of life (15-17). Most, however, have a limited phylogenetic distribution, being present in subsets of bacterial, archaeal or eukaryotic lineages. Orthologs of DnaJ of *E. coli*, have by far the broadest phylogenetic distribution. Called class A or type I JDPs, they are present in most bacterial, in some archaeal and in the vast majority of eukaryotic lineages (18, 19). In eukaryotic cells they are present in all major sub-cellular compartments: mitochondria, plastids, the endoplasmic reticulum (ER) and the cytosol/nucleus (C/N) (15). Class A JDPs are structurally complex (Fig. 1), having an N-terminal J-domain, an adjacent glycine/phenylalanine rich (G/F) region, followed by two β-sandwich substrate binding domains (CTD1 and CTD2), a dimerization domain (DD) and a C-terminal extension (CTE). All class A JDPs have a zinc finger domain (ZnF) protruding from the CTD1 – their diagnostic feature (Fig. 1) (18). The ZnF is known to be involved in substrate interactions (13, 20, 21).

**Fig. 1.**
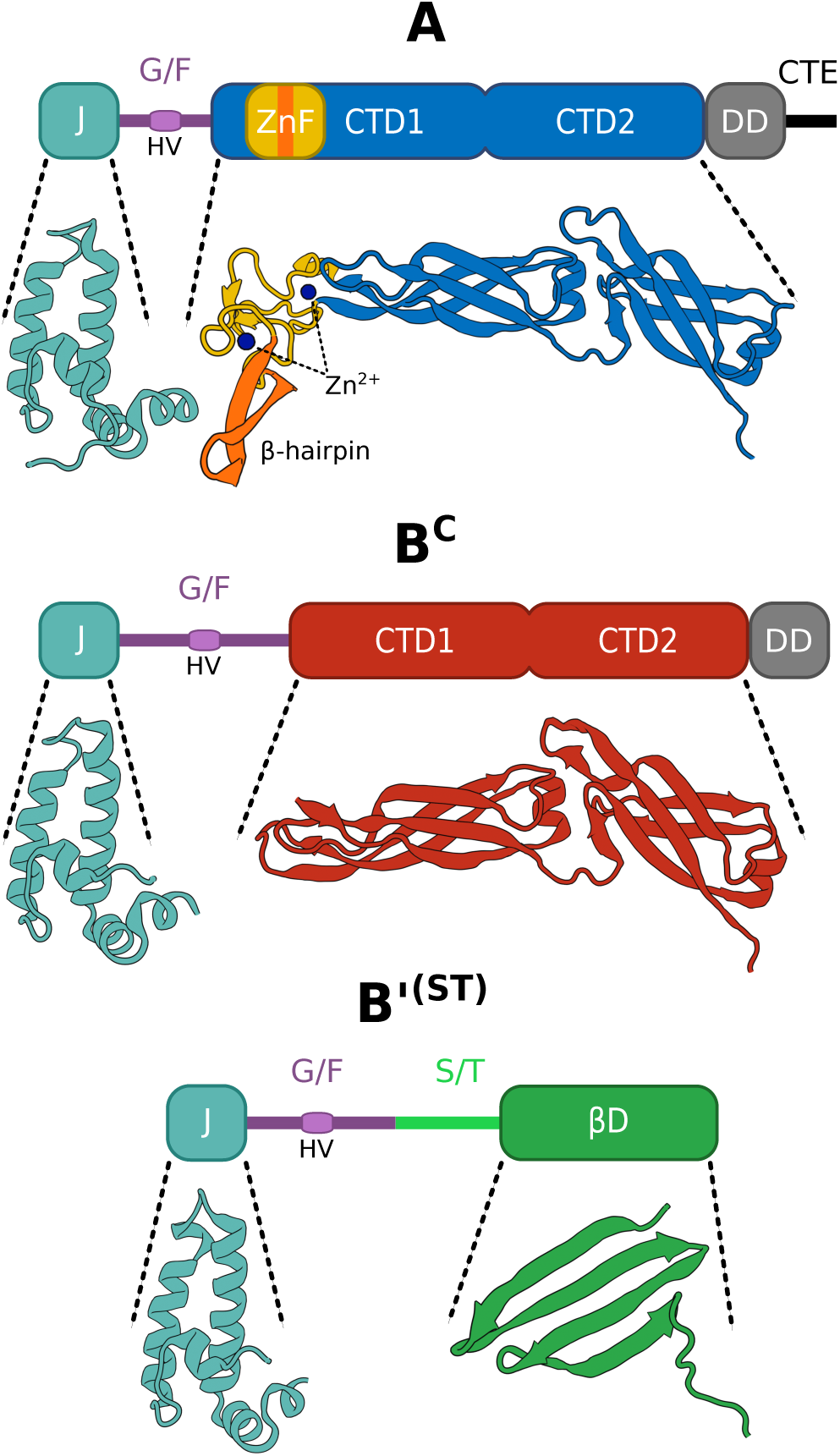
Domain organization of class A and B JDPs. Line diagrams of three classes of human JDPs: A, top; B^C^, middle; B’^(ST)^, bottom (structured domains, rectangles; unstructured regions, lines), with structures of J-domains and substrate binding domains shown. All three have an N-terminal J-domain (J; cyan) followed by a disordered glycine/phenylalanine rich region (G/F; violet), containing a short helical segment termed helix V (HV). Class A and B^C^ JDPs have two C-terminal substrate binding domains (CTD1, CTD2); blue and red, respectively and a dimerization domain (DD; grey) after the G/F. As have a defining zinc finger region (ZnF; yellow/orange) extending from CTD1, as well as a C-terminal extension (CTE; black) following the DD. B’^(ST)^s have an unstructured serine/threonine rich segment (ST; light green) followed by substrate binding β-pleated sheet domain (βD; green). Structures: DNAJA2 - J (PDB: 2lo1), CTD1/2 (PDB:7zhs); DNAJB1 - J (PDB: 6z5n), CTD1/2 (PDB:3agx); DNAJB6 – J (PDB: 6u3r), βD (PDB: 7jsq)

In both bacteria and compartments of eukaryotic cells, Class A JDPs are often accompanied by JDPs of the B class (type II). By definition, class B, like class A, JDPs have an N-terminal J-domain and an adjacent G/F region, but lack a ZnF (18). Class B JDPs can be divided into two types. Some, termed canonical B (B^C^), such as CbpA of *E. coli*, share the substrate binding CTD1/2 and dimerization domains (DD) with class A members (Fig. 1). Others, which we refer to throughout as B’ JDPs, do not have CTD1/2 or DD. The best studied B’ JDPs are those that are found in the cytosol/nucleus of metazoans and have a serine/threonine rich segment (ST) followed by a substrate binding β-pleated sheet domain (βD) immediately after the G/F region (6). These are referred to as B’^(ST)^ throughout (Fig. 1). Class A and B JDPs are involved in general chaperone functions, including protein refolding after stress (1, 7). But each, upon interacting with specific substrates, drive distinct processes. For example, class A JDPs, via their ZnF, prevent aggregation of substrates with unstable β-sheets (e.g. mutant p53 oncoprotein) (21), while Bs have been repeatedly tied to amyloids (6, 22).

Amyloids, highly ordered β-sheet-rich protein fibrils, are found in all domains of life (23-25). In some cases, amyloids play a functional role (e.g. polypeptide hormones are stored in a β-sheet conformation; bacteria use β-sheet fibrils in biofilms). In other cases, amyloids are pathogenic (e.g. prions cause Creutzfeldt-Jakob disease). In fungi B^C^ JDPs (i.e. Sis1 in *Saccharomyces cerevisiae*) are an essential part of the system that fragments many types of yeast prions enabling their passage from one generation to the next (22). In humans amyloids are associated with a variety of disease states, particularly neurodegenerative disorders including Alzheimer’s, Parkinson’s and Huntington’s diseases. Human B^C^ JDPs (i.e. DNAJB1) drive Hsp70 dependent, amyloid disaggregation, while B’^(ST)^ JDPs (i.e. DNAJB6 and B8) are particularly potent in preventing amyloid fibrils formation (6, 7, 26-28).

While recent research has provided much new information about the functional similarities and differences among JDPs from diverse systems, the fundamental question of the origin of JDPs, particularly B^C^s and B’^(ST)^s, has not been addressed. Because such information is relevant to understanding the origins of functional differences between class A and B JDPs and the ability of class B JDPs to affect amyloids, we carried out extensive phylogenetic analyses. Our results revealed that B^C^s evolved independently more than once from class A ancestors, convergently losing their ZnF. Using biochemical resurrection of inferred ancestral sequences, we found that the ability of cytonuclear class B^C^ JDPs to disassemble amyloid fibrils emerged early, but subsequent to ZnF loss.

## Results and Discussion

### Class B^C^ JDPs evolved independently more than once from class A ancestors

To resolve the evolutionary origin of class B JDPs, we carried out an extensive phylogenetic analysis using class A and B (B^C^ and B’^(ST)^) sequences retrieved from 747 bacterial, archaeal and eukaryotic genomes. Of the 1,420 retrieved sequences, 710 had a full-length ZnF, thus forming the class A group, with the remaining, having no or a truncated ZnF, forming the class B^C^ (556) and class B’^(ST)^ (154) groups. We obtained well resolved and highly congruent phylogenetic trees with strong support for major JDP groups regardless of the method/model used or the presence/absence of B’^(ST)^ sequences in a dataset (Fig. 2, suppl. Fig. S1-12). The trees revealed a complex phylogenetic distribution of class B^C^ JDPs. Not all members of this class cluster together as expected if B^C^s originated once in bacteria and were subsequently inherited by eukaryotic cells. Three B^C^ monophyletic groups branched off from class A JDPs, independently from the CbpA clade that contains most bacterial B^C^s – one in a subset of bacteria (Actinobacteria) and two in the eukaryotic lineage (Fig. 2).

**Fig. 2.**
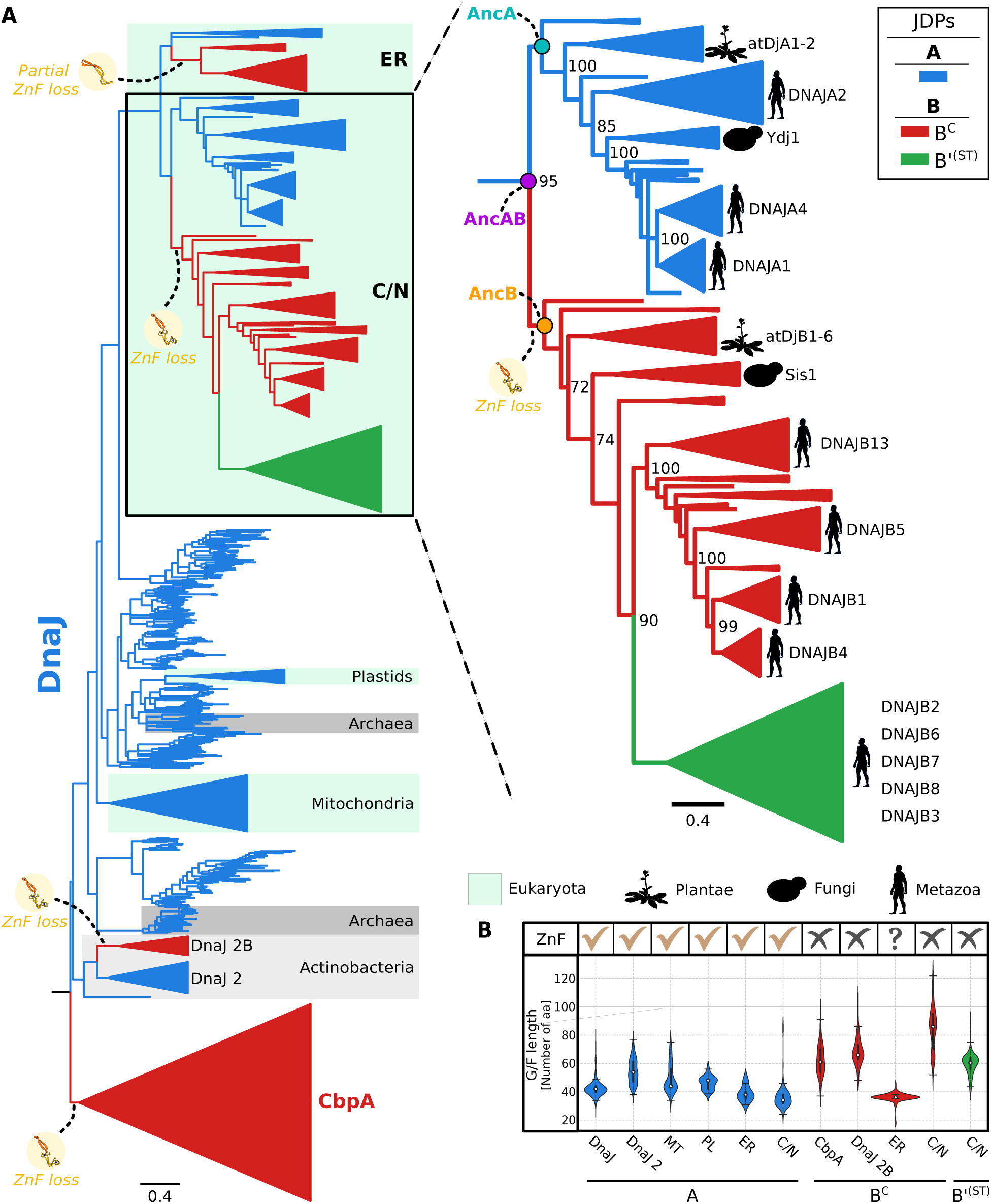
Evolutionary history of class A, B^C^ and B’^(ST)^ JDPs. (A) The best supported tree that includes class A JDPs (blue), B^C^ JDPs (red) and B’^(ST)^ JDPs (green) (suppl. Fig. S1). Left: JDPs highlighted: Archaea (dark gray); Actinobacteria (light gray, with additional JDP copies (DnaJ 2 and DnaJ 2B) indicated; eukaryotic subcellular compartments mitochondria, plastids, cytosol/nucleus (C/N), endoplasmic reticulum (ER), (turquoise). Right: Zoom in for JDPs of C/N. Clades containing metazoans, fungi and plants are collapsed into triangles. Phylogenetic position of representative members of class A, B^C^ and B’^(ST)^ JDPs from *Saccharomyces cerevisiae*, *Arabidopsis thaliana*, and *Homo sapiens* indicated. For JDPs from the C/N, circles represent the last common ancestors of class A and B^C^/B’^(ST)^ (AncAB, magenta), class A (AncA, cyan) and class B^C^/B’^(ST)^ (AncB, orange). Loss or partial loss of a ZnF region is indicated (yellow circles). Scale bar – amino acid substitutions per position. Bootstrap support values for major splits are indicated. Splits with bootstrap < 50 were collapsed into polytomy. (B) Presence of ZnF and length of the G/F region. ZnF presence (✔) or absence (✘); question mark symbolizes truncated ZnF. Violin plots represent the distribution of the number of residues constituting the G/F region. Median is marked with white dot; black boxes represent first and third quartiles (Q1 and Q3); plot width is proportional to the number of sequences with a given number of residues. DnaJ – bacterial class A, DnaJ 2/DnaJ 2B – DnaJ class A/B^C^ homologs from Actinobacteria, MT – mitochondria, PL – plastids, ER – endoplasmic reticulum, C/N – cytosol/nucleus, CbpA – bacterial B^C^.

Our phylogenetic analysis traces the origin of uncharacterized B^C^ JDPs from Actinobacteria to a DnaJ duplicate (DnaJ 2) that emerged at the stem of this bacterial lineage, followed by complete ZnF loss in a subset of Actinobacteria. In the case of cytonuclear B^C^ JDPs, which also completely lack a ZnF, their origin was traced to two sequential class A gene duplications that took place in a common ancestor of all eukaryotes (Fig. 2, 3). The first led to a common ancestor of all the A/B^C^s present in both the ER and the cytosol/nucleus. A subsequent duplication of the cytosol/nucleus A led to sister cytonuclear class A and B^C^ JDPs. Most metazoans and plants have multiple B^C^s (e.g. DNAJB1, B4, B5; B13 in humans and six, AtDjB1, B2, B3, B4, B5, B6, in *A. thaliana*). In both, the multiple copies are more closely related to each other than to any class A or to B^C^ JDP from other lineages, including the single B^C^ present in fungi (e.g. Sis1 in *S. cerevisiae*). Such similarity indicates that multiplication of cytonuclear B^C^s was driven by lineage specific duplications of B^C^ genes. For example, three sequential duplications of B^C^ genes can explain the presence of the four B^C^ paralogs in humans (Fig. 2, 3).

In sum, the branching pattern of class B^C^ JDPs indicates that each class B^C^ group shares common ancestry with a different class A JDP. More specifically, class B^C^ JDPs from the cytosol/nucleus of eukaryotic cells and from Actinobacteria are more closely related to their class A predecessor than to members of other B^C^ groups. These results imply that B^C^s independently evolved from class A ancestors via loss of the ZnF. That loss of the ZnF occurred independently, that is convergently, more than once raises the question of whether such loss enabled evolution of distinctive functional abilities, while substantially maintaining overall function. This idea is consistent with previous results (20, 21). For example, deletion of the ZnF’s β-hairpin (Fig. 1), while responsible for its unique interaction with oncogenic mutant of p53, does not affect interaction with other substrates (21). Thus, though the ZnF may have broadened substrate binding ability, both the innate rigidity of the ZnF (Fig. 1) and the inflexibility it imposes on the homodimer (29-31) may restrict conformational dynamics that are required for the unique functionality of B^C^s.

### Human cytonuclear B’^(ST)^ JDPs emerged at the base of metazoans from a B^C^ ancestor

We also conclude from our phylogenetic analysis that B’^(ST)^s originated from a cytonuclear B^C^ in a common ancestor of metazoans. On our best supported trees, B’^(ST)^s share a common ancestry with metazoan cytonuclear B^C^s and are only distantly related to fungal (Sis1) and plant (atDjB1-6) B^C^s (Fig. 2, suppl. Fig. S1-3). Furthermore, the five human B’^(ST)^ paralogs (DNAJB2, B3, B6, B7, B8) are more closely related to each other than to any class B^C^ or A JDP, indicating they evolved by sequential duplications of B’^(ST)^ genes. We compared the distribution of B’^(ST)^s across metazoans with the branching order of their phylogeny to determine evolutionary relationships amongst them (suppl. Fig. S13). We found that DNAJB2 branches off at the base of the B’^(ST)^ tree and is present in all metazoans, thus representing the predecessor of the B’^(ST)^s lineage (Fig. 3, suppl. S13). Its duplication in a common ancestor of vertebrates gave rise to the DNAJB6 lineage. DNAJB6’s subsequent duplication in a common ancestor of reptiles and mammals gave rise to the DNAJB8 lineage. Later, two independent duplications of DNAJB6 and DNAJB8 in a common ancestor of mammals gave rise to DNAJB7 and DNAJB3 respectively.

**Fig. 3.**
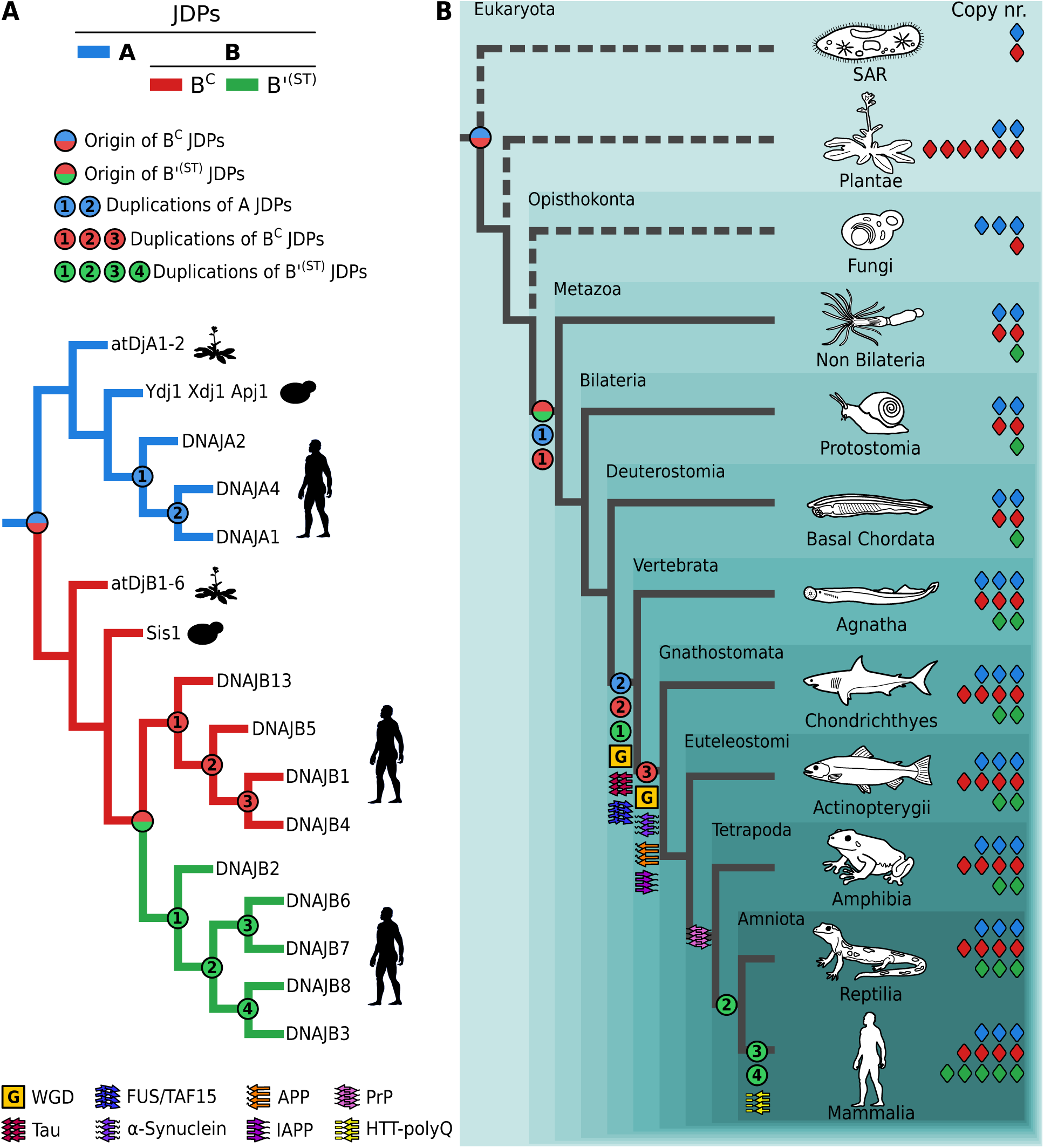
Evolutionary origin and expansion of the cytonuclear JDPs in the human lineage. Left: Simplified phylogeny of cytonuclear JDPs with names of *A. thaliana*, *S. cerevisiae* and *H. sapiens* A, B^C^ and B’^(ST)^ JDPs. Right: Placement of JDP gene duplication events and emergence of human amyloidogenic precursor proteins across simplified phylogeny of eukaryotes. Color coding of branches leading to JDP clades at left and gene number at right (diamonds): A (blue), B^C^ (red), B’^(ST)^ (green). Gene duplication events are marked with circles: blue/red - emergence of B^C^ from A ancestor. red/green - emergence of class B’^(ST)^ from B^C^ ancestor, blue (1, 2) sequential duplications of As; red (1, 2, 3) sequential duplications of B^C^s; green (1, 2, 3, 4) sequential duplications of B’^(ST)^s. Whole genome duplication events (G) and graphical symbols representing emergence of predecessors of human amyloidogenic proteins are indicated. Dashed branches lead to non-metazoan eukaryotes.

Thus, four gene duplications can explain the presence of the five B’^(ST)^ JDPs in humans (Fig. 3). We also noted that the DNAJB2 duplication coincided with a whole genome duplication (WGD) event at the base of vertebrates (Vertebrata) (32). This and a subsequent WGD event at the base of jawed vertebrates (Gnathostomata) could have also been responsible for duplications of class A and B^C^ JDPs (Fig. 3). We also note that human DNAJB3 does not have S/T and βD due to a premature stop codon. The biological significance of this is not clear (33), as it is the exception. All DNAJB3 orthologs from our dataset, including chimpanzee – the most closely related species to humans (34) – do not have a stop codon, and thus have both S/T and βD.

The ancestry of S/T and βD domains of the B’(ST) is not established by our analysis. These domains do not align with either B^C^ or A sequences. However, both βD and CTD1/2 of As and B^C^s are enriched in β-strands (Fig. 1). βD might have originated from CTD1/2 and then undergone major rearrangements, obscuring positional homology between ancestral and daughter sequences (35). We therefore carried out analyses based on pair-wise comparisons of Hidden Markov Models (HMM) sequence profiles, which capture conservation of each amino acid position in the sequence alignment, allowing detection of even weak homology (36). We prepared profiles of segments of βD and CTD1/2, as well as full J-domain and G/F profiles of B’^(ST)^, B^C^ and A JDPs (suppl. Fig. S14). Consistent with our phylogenetic results, the J-domain and G/F regions of B’s are more similar to those of B^C^s than those of As. However, no sequence homology between the βD and S/T profiles of B’^(ST)^s and those based on B^C^ and A sequences was detected, suggesting that the C-terminal domains of B’^(ST)^ did not evolve from B^C^ or A sequences. Therefore, B’^(ST)^s likely have a chimeric evolutionary origin. S/T and βD could have evolved via fusion with another gene (37). However, our B’^(ST)^ profiles did not detect any homologous sequences in prokaryotic or eukaryotic proteomes. Possibly, S/T and βD underwent such strong rearrangements during evolution that detectable homology with source sequence(s) was lost (38) or evolved *de novo* from noncoding DNA segment(s) (39).

Regardless of the origin of the S/T and βD domains, the phylogeny discussed above (Fig. 2, 3) demonstrates that the J-domain and G/F of B’^(ST)^s descended from a cytonuclear B^C^ predecessor. This origin is consistent with the shared features of the G/F regions. The G/F length of both cytonuclear B^C^s and B’^(ST)^s is longer than that of class A JDPs (Fig. 2B). Furthermore, the helical region (helix V) (Fig. 1) within the G/F of B’^(ST)^ interacts intramolecularly with its J-domain (40), regulating activity, as does helix V of class B^C^ JDPs (41, 42). We speculate that a longer G/F enables autoregulatory activity of helix V is critical for the amyloid related activities of both B^C^s and B’^(ST)^s.

### Expansion of cytonuclear B^C^ and B’^(ST)^ JDPs predates emergence of pathogenic amyloids in the human lineage

The known ability of human B’^(ST)^ and B^C^ JDPs to engage with amyloids prompted us to investigate the relationship between their emergence/proliferation and the emergence of amyloidogenic proteins in the human lineage (Fig. 3). The only disease-causing amyloidogenic protein for which we found information about emergence in the literature was the microtubule-associated protein (Tau) (43). We therefore reconstructed phylogenies of seven others for which there is experimental evidence that JDP/Hsp70 systems either affect their fibrilization or are able to dissociate their fibrils: Aβ protein precursor (APP), fused in sarcoma (FUS), TATA-box binding protein-associated factor 15 (TAF15), islet amyloid polypeptide (IAPP), α-synuclein (α-syn), prion protein (PrP), huntingtin protein (HTT) (suppl. Fig. S15-18). Our analyses revealed that Tau, as well as a FUS/TAF15 progenitor, emerged at the base of vertebrates (Vertebrata).

Others emerged later: APP, IAPP, and α-syn at the base of jawed vertebrates (Gnathostomata), PrP at the base of four-legged vertebrates (Tetrapoda) and HTT-polyQ at the base of mammals (Mammalia) (Fig. 3). Thus, the emergence of all these amyloidogenic proteins was predated by both the emergence of the B^C^ at the base of eukaryote and the emergence of B’^(ST)^ at the base of metazoans. Subsequent duplications of both B’^(ST)^ and B^C^ coincided with the emergence of Tau and FUS/TAF15 precursor, while the final duplication of B^C^ coincided with the emergence of APP, IAPP, and α-syn (Fig. 3). The three duplications of B’^(ST)^ that took place at the base of amniotes (Amniota) (one duplication) and Mammalia (two duplications) roughly coincided with the emergence of PrP and HTT-polyQ.

Taken together, our results indicate that the B′^(ST)^ subfamily arose through a B^C^-gene duplication before the appearance of the disease-causing amyloids we examined, while the subsequent expansion of cytonuclear B′^(ST)^ and B^C^ paralogs in the human lineage coincided with their emergence. Thus, JDPs that can both prevent formation and facilitate disassembly of amyloid fibrils not only were in place before emergence of amyloid forming proteins in the human lineage, but they also multiplied in their presence.

### Ancestral cytonuclear B^C^ is functionally similar to contemporary B^C^s and different from both ancestral and contemporary class A JDPs

The engagement of cytonuclear B^C^s, as well as the B′^(ST)^s that emerged from them, with amyloids, motivated us to ask whether such functional properties emerged before or after ZnF loss. We compared the biochemical properties of resurrected ancestral proteins (44) before (AncAB) and after (AncB and AncA) the class A gene duplication that led to B^C^ emergence (Fig. 2). Reconstructed AncB has a domain composition typical of present-day B^C^s, including the lack of the ZnF, while AncAB and AncA have all the domains present in contemporary class A JDPs, including a full length ZnF (suppl. Fig. S19).

To assess whether these reconstructed ancestral JDPs are functional, we tested their ability to substitute for present-day A and B^C^ in *in vivo* growth and biochemical protein refolding assays, using the budding yeast and human systems, respectively. Yeast strains having either a deletion of the chromosomal gene encoding B^C^ Sis1 or A Ydj1 (Fig. 4B) were used (45, 46). Consistent with expectations – AncB supported growth of *sis1*Δ cells as well as Sis1 itself, but not *ydj1*Δ; AncA supported robust growth of *ydj1*Δ, but only minimal growth of *sis1*Δ. AncAB did not support robust growth of either strain, albeit improved growth of *ydj1*Δ substantially more than *sis1*Δ. We also tested the ability of purified ancestral proteins to refold a denatured substrate protein, luciferase. Both human class A (DNAJA2) and B^C^ (DNAJB1) have been shown, in conjunction with human Hsc70 and nucleotide exchange factor (NEF) Hsp105, to be functional in this assay (47, 48). The three ancestral JDPs had substantial activity, as well (Fig. 4C). Based on these *in vivo* and *in vitro* results we concluded that these ancestral JDPs have activity, allowing us to productively compare their functional properties.

**Fig. 4.**
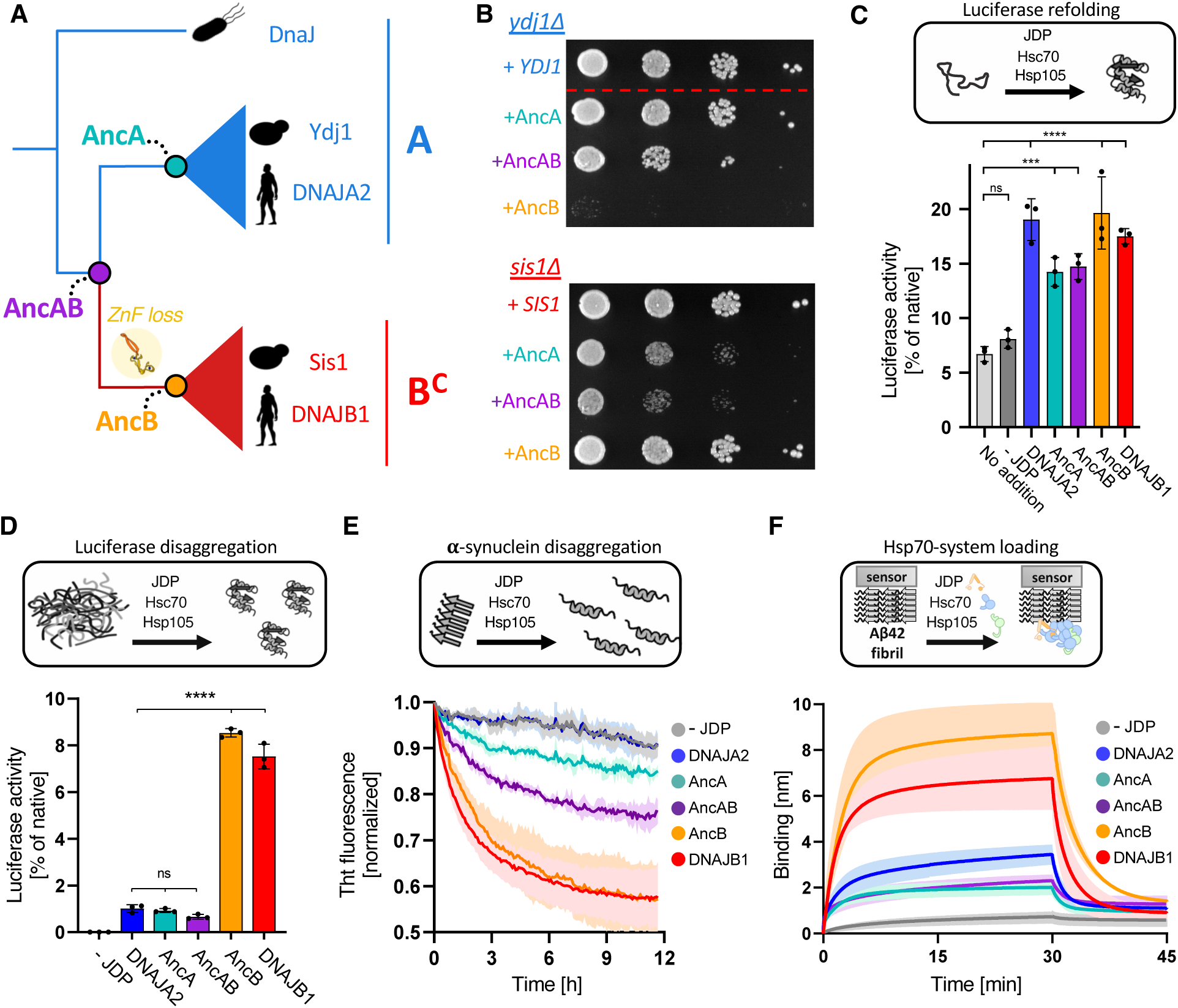
Functional divergence of B^C^ JDPs from the cytosol/nucleus. (A) B^C^s from the C/N share common ancestor (AncAB) with class A JDPs. Following AncAB gene duplication canonical Bs (B^C^) diverged from class A JDPs losing a ZnF region, indicated on the branch leading to AncB. Sequences representing the common ancestors of B^C^s (AncB) and As (AncA) as well as AncAB were reconstructed for use in subsequent experiments (B-F). (B) Ancestral JDPs are functional *in vivo* in yeast. *ydj1*Δ (top) and *sis1*Δ (bottom) cells harboring plasmid-borne copies of ancestral JDPs, WT *YDJ1* or WT *SIS1*, as indicated, were plated as 10-fold serial dilutions on glucose-rich medium and incubated at 30°C for 2 days. Dotted line indicates removal of nonrelevant sample from image. (C) Purified ancestral JDPs are active in chaperone-assisted refolding of Fire-fly Luciferase (FLuc). FLuc refolding at 200 nM was assayed upon dilution from 5M guanidinium-HCl into buffer without (spontaneous refolding) or with chaperones: Hsc70 at 3 μM, Hsp105 at 0.3 μM and either contemporary or ancestral JDPs at 1 μM, as indicated. FLuc activity was normalized to that of native protein; average activity of at least 3 repeats ± standard deviation is shown. (D) AncB, but neither AncAB nor AncA, is active in FLuc disaggregation. Disaggregation of FLuc aggregates at 200 nM in the presence of Hsc70 at 1.5 µM, Hsp105 at 0.15 µM and JDPs at 1 µM, as indicated. FLuc activity was normalized to that of native FLuc; average activity of at least 3 repeats ± standard deviation is shown. (C), (D) Two-way ANOVA: **p>0.01, ***p>0.001, ****p>0.0001. (E) AncB is more active than either AncAB or AncA in α-synuclein fibrils disassembly. Disassembly of α-synuclein fibrils in the presence of Hsc70 at 3 µM and Hsp105 at 0.3 µM, and JDPs at 0.25 µM, as indicated. Fibril disassembly was monitored by fluorescence of ThT, an amyloid-specific dye that reports on the total amyloid load in the sample. (F) AncB drives interaction of Hsc70 and Hsp105 with Aβ42 amyloid fibrils more efficiently than either AncAB or AncA. Interaction of Hsc70 at 1 µM and Hsp105 at 0.1 µM with Aβ42 amyloid fibrils immobilized to a BLI biosensor was monitored in the absence or presence of JDPs at 1 µM. (E), (F) The lines represent the average of three replicates, with shading designating standard deviation.

Next, we tested ancestral JDPs in biochemical assays in which present-day canonical B^C^s are more active than As: (i) disaggregation of amorphous aggregates, (ii) disassembly of amyloid fibrils, (iii) recruitment of Hsp70/NEF to amyloid fibrils. Indeed, in disaggregation of pre-formed luciferase aggregates, carried out in the presence of Hsp70/NEF, AncB was as efficient as present-day B^C^s, that is human DNAJB1 or yeast Sis1 (Fig. 4D, suppl. Fig. S20). AncA and AncAB had marginal activity comparable to DNAJA2. We observed a similar pattern in the assay, in which disassembly of pre-formed α-synuclein fibrils (suppl. Fig. S21) in the presence of ancestral proteins and human Hsp70/NEF was assessed (Fig. 4E, suppl. Fig. S22). AncB was more active than either AncAB or AncA. The recruitment assay measured the ability of JDPs to facilitate interaction of Hsp70/NEF with pre-formed Aβ42 amyloid fibrils immobilized to an optical sensor (suppl. Fig. S23). AncB drove fast binding kinetics and high binding yield, similar to DNAJB1, while binding signals for both AncA and AncAB were low, comparable to DNAJA2 (Fig. 4F, suppl. Fig. S24). We interpreted these results as consistent with our hypothesis that AncB diverged functionally, gaining novel biochemical activities characteristic for present day B^C^s, subsequent to its emergence via a class A gene duplication.

Taken together, our experimental results indicate that AncB is more efficient than AncAB and AncA in recruiting Hsp70/NEF chaperones to protein aggregates and in disassembling them. Therefore, we concluded that novel activities that differentiate cytonuclear B^C^s from class A JDPs evolved subsequent to ZnF loss – early on the branch connecting AncAB with the last common ancestor of the B^C^ lineage (AncB) (Fig. 4A).

### Class B^C^ JDPs of the ER emerged from class A by partial loss of the ZnF

Evolutionary relationships, as well as the basic classification, of class A/B^C^ JDPs of the ER has been debated for years based on information from model systems. Fungi (i.e. Scj1 in *S. cerevisiae*) have a class A JDP; metazoans and plants (i.e. DNAB11 and atDjA8 in human and *Arabidopsis thaliana*, respectively) have a JDP with a truncated ZnF - 30 rather than the 60 amino acids (with 4, not 8 cysteines) found in full-length Class A ZnFs (17, 49, 50). Analysis of our dataset demonstrated the broad presence of a full-length ZnF in fungi and unicellular eukaryotes, while a truncated ZnF is present in metazoans and plants (suppl. Fig. S25). All belong to a single clade (Fig. 2, 5) - with fungal As only distantly related to metazoan and plant B^C^s - inconsistent with the well-established sister relationship of metazoans and fungi, and their distant relationship to plants (Fig. 5) (51). Emergence of class B^C^ JDPs after a class A gene duplication in a common ancestor of metazoans, fungi and plants (Fig. 5) could explain our observed relatedness between ER JDPs - if one duplicate subsequently evolved a truncated ZnF (class B^C^), while the other maintained a full-length ZnF (class A), followed by differential gene loss (Fig. 5).

**Fig. 5.**
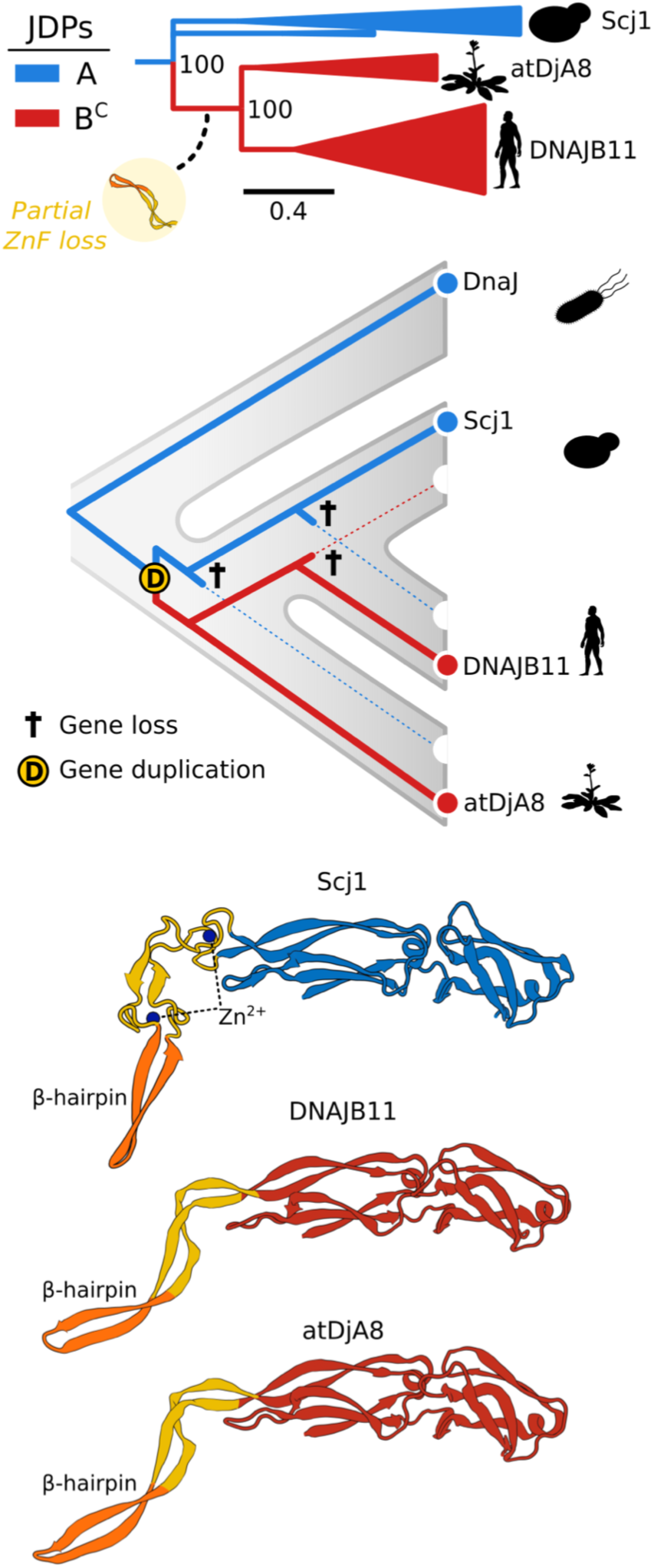
Evolution of class A and B^C^ JDPs from the ER. Top: Phylogeny of class A and B^C^ JDPs from the ER - class A (blue) and B^C^ (red) with names of *A. thaliana*, *S. cerevisiae* and *H. sapiens* JDPs. Clades containing metazoans, fungi and plants are collapsed into triangles. Partial loss of ZnF (yellow circle). Scale bar – amino acid substitutions per position. Bootstrap support for major splits is indicated. Middle: Scenario that could explain the emergence of B^C^s with a shorter ZnF and the evolutionary relationships among class A (blue) and B^C^ (red) JDPs from the ER presented in the context of species phylogeny (grey). “D” marks duplication event prior to ZnF shortening (red line); cross (†) indicates the loss of a gene copy in a particular evolutionary lineage. Bottom: AlphaFold structural models of three contemporary JDPs of the ER: yeast Scj1 (ID:AF-P25303-F1), human DNAJB11 (ID: AF-Q9UBS4-F1) *A. thalina* atDjA8 (ID: AF-Q9LZK5-F1). Full length ZnF of Scj1 and truncated ZnF of DNAJB11 and atDjA8 (yellow) and ZnF’s β-hairpin (orange).

The longstanding discussion about ER JDPs has focused on whether plant/metazoan ER B^C^s are functionally divergent from the As of fungi (15, 52). We followed the strict definition of class A JDPs (18) and placed the ER JDPs with a truncated ZnF in class B^C^. However, similar structures are predicted for short and full length ZnF regions (50) with the β-hairpin critical for substrate interaction (21) similarly protruding (Fig. 5). Considering the oxidative environment of the ER, a short ZnF stabilized by disulfide bonds (49) could result in a structure similar to that of zinc coordinated full-length ZnF of *E. coli* DnaJ and cytosolic *S. cerevisiae* Ydj1 (29, 30), and thereby maintaining Class A functionality.

The 36-residue median length of the G/F region of B^C^s of the ER is also consistent with class A-like functionality (Fig. 2B). This length is very similar to that of As, not only of the ER, but of other eucaryotic compartments and bacteria, whose median length ranges from 34 to 54. In contrast, the median G/F region length of cytonuclear A and B^C^ JDPs is 34 and 86, respectively. Although length differences have been noted for years for As and B^C^s of yeast and mammalian model systems, the pervasiveness of the difference across species is notable. This difference may be in part attributable to the specialized inhibitory mechanism in which the helical region (helix V) in the G/F folds back on the J-domain regulating its activity (40-42). However, although not as great, the G/F of CbpA and DnaJ 2B of Actinobacteria are also longer (on average) than those of A JDPs. This increase in length must have evolved convergently, implying that it is important for their functional divergence from class A antecedents as B^C^s originated independently from class A genes.

## Conclusions

Broad taxonomic distribution (15, 16) of class A and B^C^ JDPs has long been assumed to reflect their deep evolutionary origin at the stem of the bacteria lineage. But our detailed phylogenetic analyses revealed that class B^C^ JDPs emerged independently more than once from class A ancestors - in bacteria and in eukaryotes. Nevertheless, our findings do lend support to the traditional, DnaJ centered, classification of JDPs (18). Both our results and recent surveys of JDP diversity (15, 16) indicate that class A members have the broadest phylogenetic distribution. Thus, DnaJs are likely the progenitor of all JDPs, even those sharing only a J-domain with class A and B members (i.e. class C (5)), receiving their J-domains from DnaJ homologs, as recently proposed for the class C JDP that function in the biogenesis of iron-sulfur clusters, HscB (53).

That loss of the ZnF in a Class A progenitor occurred independently, that is convergently, in bacteria and eukaryotes suggests that this structural simplification allowed acquisition of new functions and can be considered an example of reductive evolution at the protein level. Such domain losses during evolution of eukaryotic protein families is common (54) and thought to have contributed to accelerated evolution of novel gene paralogs (55-57). Once in place, both B^C^s and B’^(ST)^s duplicated several times. The increase in gene number of both B^C^ and B’^(ST)^s is consistent with a recent proposal that the demand for chaperone activity upon expansion of the proteome across the tree of life (58) was often fulfilled, not by the emergence of new chaperone families, but by an expanded network of co-chaperones, including JDPs (59). We speculate that there is such a causal relationship between multiplication of pathogenic amyloids and the presence of class B JDPs in the cytosol/nucleus of human lineage. Moreover, the presence of B^C^s and B’^(ST)^s able to effect amyloid transactions prior to the emergence of potentially pathogenic amyloidogenic proteins in the human lineage may well be an example of preadaptation – that is, did not arise by selection for their current role (60).

## Materials and methods

### Dataset of class A and class B JDPs for evolutionary analyses

To resolve evolutionary relationships among A, B^C^ and B’^(ST)^ JDPs we assembled a dataset of 747 proteomes from bacteria, archaea, plants, fungi, metazoans and protists We retrieved JDP sequences using Hidden Markov Model (HMM) sequence profiles. We used Hmmer1.3b1. to generate 2 profiles based on the well characterized class A and B^C^ sequences from model systems (*E. coli*, *A. thaliana*, *S. cerevisiae* and *H. sapiens*) aligned with Clustal Omega v1.2.2. The first profile encompassed J-domains and G/F regions (JD-GF-HMM) – diagnostic features of class A and B^C^/B’ JDPs. The second profile encompassed C-terminal domains (CTD-DD-HMM) – diagnostic feature of class A and B^C^, but not B’. Sequences retrieved by both profiles that also had a ZnF were assigned to class A, those that do not have a full length ZnF, were assigned to class B^C^. Sequences retrieved only by JD-GF-HMM but not by CTD-DD-HMM were assigned as class B’. Class A and B^C^ sequences were aligned and curated manually – incomplete or unusually fast evolving sequences were removed. Following realignment of curated A and B^C^ sequences, a full length HMM profile was built (AB^C^-HMM). This profile was used to prepare two alignments of A/B^C^ sequences for phylogenetic analyses: one for all A/B^C^ sequences from our dataset (AB^C^-alignment); the other restricted to eukaryotic A/B^C^ sequences (AB^C^E - alignment). To resolve the evolutionary origin of cytonuclear B’^(ST)^ JDPs of metazoans, whose diagnostic features are a S/T region followed by a βD domain, we retrieved B’^(ST)^ sequences from our dataset of B’ JDPs using the B’^(ST)^-HMM profile based on the alignment of four human B’^(ST)^s: DNAJB2, B6, B7 and B8. For phylogenetic analyses these sequences were aligned to the AB^C^-HMM profile along with class A and B^C^ sequences; AB^C^B’^(ST)^ -alignment. To analyze evolutionary relationships among B’^(ST)^, the B’^(ST)^ dataset was completed by addition of sequences from Agnatha and basal Chordata retrieved from KEGG and OMA databases. Next the incomplete or unusually fast evolving sequences were removed resulting in B’^(ST)^ -alignment of 126 sequences. For each position of the AB^C^, AB^C^E, AB^C^B’^(ST)^, and B’^(ST)^ alignments, posterior probability, which represents a degree of confidence in each position (residue or gap), was computed using Forward-backward algorithm (61). Positions with posterior probability >0.5 were used for phylogenetic tree reconstructions (62).

### Phylogenetic tree reconstruction

Based on the AB^C^ and AB^C^B’^(ST)^ - alignments Maximum Likelihood (ML) trees were reconstructed. 1,000 ML searches (using iQtree 2.3.6) were conducted with 100 rapid bootstrap replicates using four method/model combinations: (i) LG model with proportion of invariable sites and gamma distribution (LG + I + G) (63)(ii) C10 and (iii) C20 heterogeneous mixture models (64) with the LG + G (iv) jackknife method, which provided an alternative estimate of the support for the major splits. 100 jackknife replicates were assembled by sampling randomly 60% of positions from a sequence alignment. For each sample 1,000 ML searches were conducted using the LG + I + G model. The 100 consensus trees reconstructed in this way were collated and a master consensus tree produced. The trees were rooted at the common ancestor of bacterial DnaJ (class A) and CbpA (class B^C^). We chose this rooting, because their presence in all bacterial taxonomic groups, but very limited presence (DnaJ) and absence (CpbA) in archaea, implies common ancestry in bacteria. This rooting was statistically supported when compared to alternatively rooted topologies using the likelihood-ratio test (Table S1).

For AB^C^E - alignment phylogenetic trees were also reconstructed using Bayesian approach with PhyloBayes 3.3e. We used four models of sequence evolution: (v) LG+I +G, (vi) WLSR5 (vii) C60+LG+G and (viii) CAT+LG+G. For each method/model two independent runs were performed with a total length of >4000 cycles, until the maxdiff was < 0.3 and the minimum effective size was >50. To construct the tree, the first 600 cycles were discarded as burn-in, and the topology and posterior consensus support was computed based on the trees from remaining cycles.

For B’^(ST)^ alignment the best fit JTT+I+G model was used. To obtain ML B’^(ST)^ phylogeny concordant with the species phylogeny and phylogenetic distribution of B’^(ST)^s across metazoans (suppl. Fig. S13A) the tree was constrained to enforce monophyly of a clade encompassing DNAJB8/B6/B3 homologs from mammals and reptiles (suppl. Fig. S13B). Constrained topology has a higher score than unconstrained topology using the approximately unbiased (AU) test (65). Pictures of representative members of taxonomic groups are original or modified pictures from PhyloPic (*PhyloPic: A Library of Silhouettes for Phylogenetic and Taxonomic Research.*).

### Ancestral sequences reconstruction

Ancestral (AncA, AncB and AncAB) sequences were reconstructed based on an AB^C^ alignment and phylogeny (suppl. Fig. S5). Both the alignment and the tree were pruned to reduce the number of taxa but maintain species diversity in major JDP clades. Marginal reconstruction of ancestral sequences was performed with FastML using empirical Bayes method with ML reconstruction of insertions and deletions (66). Ancestral amino acid sequences were converted into DNA sequences using codons optimal for expression in *E. coli*. These DNA sequences were cloned into plasmid pET-24a(+) for expression in *E. coli* and subsequently subcloned as BamHI-XhoI DNA fragments into similarly digested p414TEF plasmid for testing their *in vivo* functionality in *S. cerevisiae*.

### Pairwise profile-profile comparison between B’^(ST)^ and A/B^C^ domains

To compare similarity between domains of the B’^(ST)^ JDPs and domains of A and B^C^ JDPs we prepared segment-specific HMM profiles based on the A, B^C^ and B’^(ST)^ sequences from metazoans. Sequences were aligned and divided into segments corresponding to structural regions (suppl. Fig. S14). HMM profiles were generated for each of these segments using the hhmake tool from HH-suite v3.3.0. Segment-specific profiles were compared in pairs (B’^(ST)^s profiles against profiles of either As or B^C^s) using the hhalign tool from the HH-suite v3.3.0 to assess the sequence similarity between corresponding domains from B’^(ST)^ and A or B^C^ JDPs.

### Phylogenetic analyses of amyloidogenic proteins

For phylogenetic analyses of amyloidogenic proteins (α**-**syn, APP, IAPP, FUS/TAF15, PrP and HTT-PolyQ), we prepared datasets of their amino acid sequences retrieved from the respective orthologous groups (HOG or OMA groups) available in the OMA database. To identify their homologs not captured by orthologous group assignment, the human amyloidogenic proteins listed above were used as query in BLASTP searches against representative proteomes from major metazoan lineages. For further validation, HMM profiles were prepared using HMMER v3.3.2 based on OMA-derived sequence sets for each protein family. These profiles were then used to search our dataset of 747 proteomes. In each case, sets were supplemented with additional sequences to provide representation of every major metazoan lineage. Final sets contained: 545 (APP), 59 (HTT), 348 (α-syn), 209 (PrP), 426 (FUS/TAF15), and 514 (IAPP) sequences, respectively. Sequences from each dataset were aligned with Clustal Omega v1.2.2. Poorly aligned positions were manually removed from the alignments. Phylogenetic trees were reconstructed using iQtree 2.3.6. For each set 1,000 ML searches were conducted with 100 rapid bootstrap replicates. The best fit sequence evolution model was selected for each dataset (see suppl. Fig. S15-18). Using these phylogenetic trees, we traced the origin of each amyloidogenic protein to a common ancestor of a clade that includes its human precursor protein (e.g. origin of α-syn was traced to the common ancestor of jawed vertebrates (Gnathostomata) suppl. Fig. S15B). In the case of HTT-PolyQ, we traced its origin to a common ancestor of mammals, as only mammalian HTTs have an expanded number of Q residues (suppl. Fig. S18).

### Identification of molecular features of class A, B^C^ and B’^(ST)^ JDPs

The presence of ZnF was recognized based on the conserved sequence block in the AB^C^B’^(ST)^ alignment corresponding to the ZnF region of bacterial DnaJ and the presence within it of 8 cysteine residues positioned to be able to coordinate two zinc ions. The length of the G/F region was calculated using AB^C^B’^(ST)^ alignment. It corresponds to the number of amino acids within the conserved sequence block matching the G/F region of bacterial DnaJ.

### *In vivo* assay in yeast

To test *in vivo* function of AncA, AncB and AncAB *S. cerevisiae* haploid strains isogenic to the W303 background and carrying a deletion of the chromosomal copy of *SIS1* or *YDJ1* (i.e., *sis1*Δ or *ydj1*Δ) were used. *sis1*Δ and *ydj1*Δ carrying a *URA3*-based plasmid having a wild-type (WT) copy of Sis1 (YCp50-*SIS1*(45)) or Ydj1 (pRS316-*YDJ1* (46)), respectively, were transformed with a plasmid encoding a test protein or, as controls, WT Sis1 or WT Ydj1, and then plated on complete minimal media plates containing 5 fluoroorotic acid (5-FOA) (TorontoResearch Chemicals Inc., Canada) (67) to select for cells that had lost the *URA3* plasmid containing the WT gene. Growth assays of the resulting strains were performed by serially diluting cells in 10-fold increments and spotting on rich media plates (YPD, [1% yeast extract, 2% peptone] (Difco Laboratories, Detroit, MI), 2% dextrose) starting with 5000 cells in the first position.

### Purification of proteins used in biochemical assays

Published protocols were used for purification of DNAJA2, DNAJB1, Hsc70, Hsp105 (47), Sis1 (68), Ydj1 (69) Ssa1, Sse1 (70) and His-tagged luciferase (71). AncA, AncB and AncAB were purified using the protocol for DNAJB1(47). α-synuclein was expressed from the pET21a-α-synuclein plasmid (a gift from prof. Bernd Bukau, University of Heidelberg) in BL21(DE3) cells at 37 °C. After cell lysis in a buffer containing 10 mM Tris-HCl pH 7.5 and 1 mM EDTA, the lysate was loaded onto Q-Sepharose (GE Healthcare) and eluted with a NaCl gradient up to 750 mM. The protein was dialysed against 5 mM KPi, pH 7.4 and then loaded onto Hydroxyapatite (BioRad) and eluted with gradient of 5-450 mM KPi, pH 7.4, followed by gel filtration on a Superdex 20 column (GE Healthcare) in buffer containing 20 mM Hepes-KOH pH 7.4. Finally, the protein was filtered through a 50 000 MWCO centrifugal filter (Millipore) to remove the high molecular mass impurities.

### Amyloid fibrils formation and imagining

α-synuclein fibrils were prepared as described (72), with modifications. 200 µM α-synuclein monomers were incubated for 15 days in buffer 20 mM Hepes-KOH pH 7.4 and 100 mM NaCl, at 37 °C, with shaking at 1,000 rpm. Fibrils were separated from monomers by centrifugation (20 000 g, 30 min). The fibrils were resuspended in 20 mM Hepes-KOH pH 7.4, sonicated and stored at -80 °C. Fibril formation was verified with ThT fluorescence and atomic force microscopy (AFM) imaging (suppl. Fig. S21).

Aβ42 and C-terminal Biotin-labeled Aβ42 peptides were synthesized via solid-phase peptide synthesis. Peptides were dissolved and monomeric fraction were isolated as described (73). To induce fibrillation 10 µM of Biotin-Aβ42 and Aβ42 peptides were mixed at 1:10 ratio in buffer 50 mM Tris-HCl (pH 7.5), 150 mM KCl, 15 mM MgCl_2_ and incubated at 37 °C for 1 hour with shaking at 750 rpm. Aβ42 amyloid fibril formation was confirmed using AFM imaging (suppl. Fig. S23).

AFM visualizations of Aβ42 and α-synuclein fibrils, were carried out using BioScope Resolve (Bruker, Bremen, Germany) at 23°C in air. Each sample was deposited onto a freshly cleaned mica surface and incubated for 5 min at room temperature, followed by washing with deionized water and drying with streams of nitrogen gas prior to AFM imaging. The ScanAsyst-Fluid+ probe (Bruker) was used for AFM imaging (resonant frequency f0 = 150 kHz; spring constant k = 0.7 N/m). Images were taken at 512 × 512 pixels with a PeakForce Tapping frequency of 1 kHz and an amplitude of 150 nm. Height sensor signal was used to display the protein image using NanoScope Analysis v1.9 (Bruker, Bremen, Germany).

### Luciferase refolding and disaggregation

Fire-fly luciferase (FLuc) refolding was carried out as described (74). Luciferase (20 µM) was chemically denatured in 5M GuHCl and 10mM DTT for 1 hour at 25 °C. The refolding reaction was initiated by 200-fold dilution into buffer (25 mM Hepes-KOH pH 7.5, 100 mM KCl, 10 mM Mg(OAc)_2_ 2 mM DTT, 5 mM ATP, 0.05% Tween20) without or with chaperones; Hsc70, 3 µM; Hsp105, 0.3 µM and JDP, 1 µM (DNAJB1, AncB, AncAB, AncA, DNAJA2).

FLuc disaggregation was carried out as described (75). FLuc (20 µM) was incubated in buffer 25 mM HEPES-KOH pH 7.5, 75 mM KCl, 15 mM MgCl_2_ with 6 M urea at 25 °C for 10 minutes. Then, it was transferred to 48 °C for 10 minutes and subsequently 25-fold diluted into buffer without urea. After 5 minutes of incubation at 25 °C, the disaggregation reactions were initiated by the addition of chaperones, at the following concentrations: Hsc70, 1.5 µM; Hsp105, 0.15 µM and JDP 1 µM (DNAJB1, DNAJA2, AncB, AncAB or AncA). FLuc and its substrate were purchased from Promega (E1702). Luciferase activity was measured using the Luciferase Assay Kit (E1501, Promega) with a GloMax® 20/20 Luminometer (Promega).

### α-synuclein disaggregation assay

Disaggregation of α-synuclein fibrils by chaperones was monitored using changes in ThT fluorescence over a 12–16 hours period as described (72). Reaction mixtures were prepared in a 96 Well Half-Area Non-Binding Microplate (Corning 3881). Preformed α-synuclein fibrils (0.8 µM) were incubated at 30 °C in buffer (25 mM HEPES-KOH (pH 7.5), 75 mM KCl, 15 mM MgCl_2,_ 2 mM DTT, 2 mM ATP, ATP regeneration system (8 mM PEP and 20 ng/µl pyruvate kinase) and 30 µM ThT) with Hsc70, 3 µM; Hsp105, 0.3 µM and JDP (DNAJB1, AncB, AncAB, AncA, DNAJA2), at indicated concentrations. Before measurement, the plate containing the completed reactions was incubated for 15 minutes at 30 °C. ThT fluorescence was measured using Tecan Spark plate-reader (excitation: 440 nm, emission: 480 nm). Background ThT fluorescence of buffer and chaperones was subtracted and all intensities normalized to the fluorescence intensity at *t* = 0 min. Data shown is the average value of three independent experiments with shaded areas representing standard deviation.

### Recruitment of Hsc70/Hsp105 to the sensor immobilized Aβ42 amyloid fibrils

To measure chaperone recruitment the bio-layer interferometry (BLI) sensors with immobilized Aβ42 fibrils or monomers, were immersed for 1800 s into solutions containing BLI buffer (50 mM Tris-HCl (pH 7.5), 150 mM KCl, 15 mM MgCl_2_) supplemented with 2 mM DTT and 5 mM ATP that contained the following chaperones: Hsc70, 1 µM; Hsp105, 0.1 µM; and JDP, 1 µM (DNAJB1, AncB, AncAB, AncA or DNAJA2). To measure chaperone dissociation, the sensors were subsequently immersed for 900 s into protein-free solution. Aβ42 fibrils or Aβ42 monomers (His-SUMO Aβ42), were immobilized for 15 min in BLI buffer onto the Streptavidin and NiNTA sensors (Octet^®^ SA and NTA Biosensors), respectively. Prior to recruitment measurements the fibrillar nature of immobilized Aβ42 was tested using anti-amyloid antibodies (Sigma-Aldrich, AB2287) at 1:100 dilution (suppl. Fig. S23). Experiments were conducted using BLItz (Pall ForteBio) and Octed K2 instruments (Satorius), operating at room temperature.

Levels of Hsc70 interacting with Aβ42 amyloid fibrils were quantified using fluorescently labelled Hsc70 (A488*-Hsc70) (suppl. Fig. S24H). Sensor with immobilized Aβ42 fibril was incubated for 30 minutes with a mixture of A488*-Hsc70 1 µM, Hsp105 0.1 µM and ancestral or contemporary JDP 1 µM in the BLI buffer - association step. Fluorescence was measured after sensor was incubated for 15 minutes in the BLI buffer without proteins - dissociation step.

## Acknowledgements and funding sources

This study was supported by National Science Center, Poland (OPUS 21 2021/41/B/NZ8/02835) (to BT). The work of A.K. and H.W. was supported by a National Science Center, Poland grant (OPUS 27; 2024/53/B/NZ1/00632). We acknowledge the Polish high-performance computing infrastructure PLGrid for awarding this project access to the LUMI supercomputer, owned by the EuroHPC Joint Undertaking, hosted by CSC (Finland) and the LUMI consortium through PLL/2023/05/016759.

