## Supplementary information for "Origin of Class B J-domain proteins involved in amyloid transactions"

**Table S1.** Likelihood values and statistical support for alternative rooting of the ML phylogeny of class A and $B^C$ JDPs.

| Outgroup | logL | ΔL | WKH | WSH | AU |
|---|---|---|---|---|---|
| CbpA | -362375.91 | 0.00 | 0.542 | 1 | 0.533 |
| DnaJ 2 | -362375.91 | $3.88*10^{-5}$ | 0.458 | 0.458 | 0.467 |
| DnaJ Archea | -362375.91 | $3.88*10^{-5}$ | 0.458 | 0.458 | 0.468 |
| CbpA+DnaJ 2 | -362375.91 | $3.88*10^{-5}$ | 0.458 | 0.458 | 0.446 |
| DnaJ+CbpA+DnaJ 2 | -362375.91 | $3.88*10^{-5}$ | 0.458 | 0.458 | 0.334 |

**logL**– log-likelihood value

**ΔL**– difference in log-likelihood compared to the best-fitting topology

**WKH**– Kishino–Hasegawa test with weights

**WSH**– Shimodaira–Hasegawa test with weights

**AU**– AU test (Approximate Unbiased test)

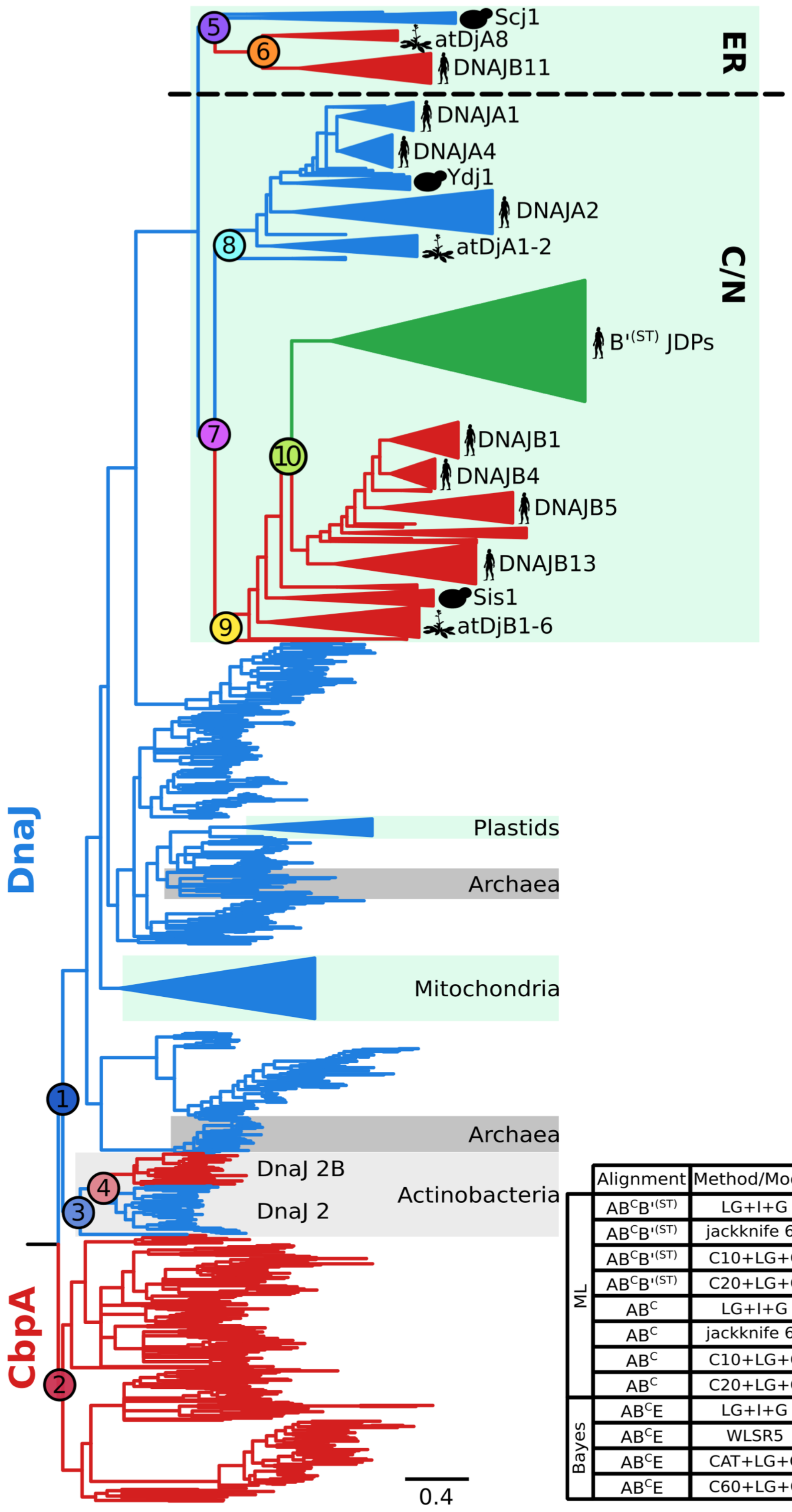


| | Alignment | Method/Model | Support for major spilts | | | | | | | | | |
|---|---|---|---|---|---|---|---|---|---|---|---|---|
| | | | 1 | 2 | 3 | 4 | 5 | 6 | 7 | 8 | 9 | 10 |
| ML | $AB^CB'^{(ST)}$ | LG+I+G | 74 | 74 | 98 | 99 | 100 | 100 | 95 | 92 | 99 | 90 |
| | $AB^CB'^{(ST)}$ | jackknife 60 | 98 | 98 | 99 | 97 | 100 | 100 | 99 | 81 | 98 | 84 |
| | $AB^CB'^{(ST)}$ | C10+LG+G | 92 | 92 | 100 | 100 | 100 | 100 | 100 | 99 | 100 | 74 |
| | $AB^CB'^{(ST)}$ | C20+LG+G | 94 | 94 | 100 | 100 | 100 | 100 | 100 | 96 | 100 | p |
| | $AB^C$ | LG+I+G | 95 | 95 | 98 | 100 | 100 | 100 | 99 | 88 | 98 | - |
| | $AB^C$ | jackknife 60 | 80 | 80 | 99 | 100 | 100 | 100 | 99 | 96 | 100 | - |
| | $AB^C$ | C10+LG+G | 88 | 88 | 99 | 100 | 100 | 100 | 100 | 99 | 100 | - |
| | $AB^C$ | C20+LG+G | 89 | 89 | 100 | 100 | 100 | 100 | 100 | 100 | 100 | - |
| Bayes | $AB^CE$ | LG+I+G | - | - | - | - | 72 | 100 | 72 | 97 | 51 | - |
| | $AB^CE$ | WLSR5 | - | - | - | - | 89 | 100 | 89 | 94 | 100 | - |
| | $AB^CE$ | CAT+LG+G | - | - | - | - | 66 | 100 | 66 | 88 | 61 | - |
| | $AB^CE$ | C60+LG+G | - | - | - | - | 75 | 100 | 75 | 100 | 58 | - |

**Figure S1.** (left) Maximum Likelihood (ML) phylogeny of A, $B^C$ and $B'^{(ST)}$ JDPs ($AB^CB'^{(ST)}$ -alignment) reconstructed with LG+I+G model. Class A JDPs (blue), class $B^C$ JDPs (red) class $B'^{(ST)}$ JDPs (green). JDPs highlighted Archaea (dark grey), Actinobacteria (light gray, with additional JDP copies (DnaJ2-B and DnaJ2-A)) indicated; eukaryotic subcellular compartments mitochondria, plastids, endoplasmic reticulum (ER), cytosol/nucleus (C/N) (light green). Dashed line - division between ER and C/N JDPs. Phylogenetic positions of representative members of class A, $B^C$ $B'^{(ST)}$ JDPs from *Saccharomyces cerevisiae, Arabidopsis thaliana, and Homo sapiens* are indicated. Nodes (circles) representing major splits are indicated. Scale bar – amino acid substitutions per position. (right) Support for major splits by different phylogenetic methods: (ML - bootstrap, Bayes- posterior probability, and jackknife) alignments and models of sequence evolution, as indicated. ML phylogenies of 1420 A, $B^C$ and $B'^{(ST)}$JDPs from bacteria, archaea and eukaryote ($AB^C$ $B'^{(ST)}$-alignment). ML phylogenies of 1266 JDPs without fast evolving $B'^{(ST)}$JDPs ($AB^C$-alignment). Bayesian phylogenies (Bayes) of 304 A and $B^C$ JDPs from the ER and C/N of eukaryote ($A/B^CE$-alignment). Tree topologies are shown on figures S2-S12.

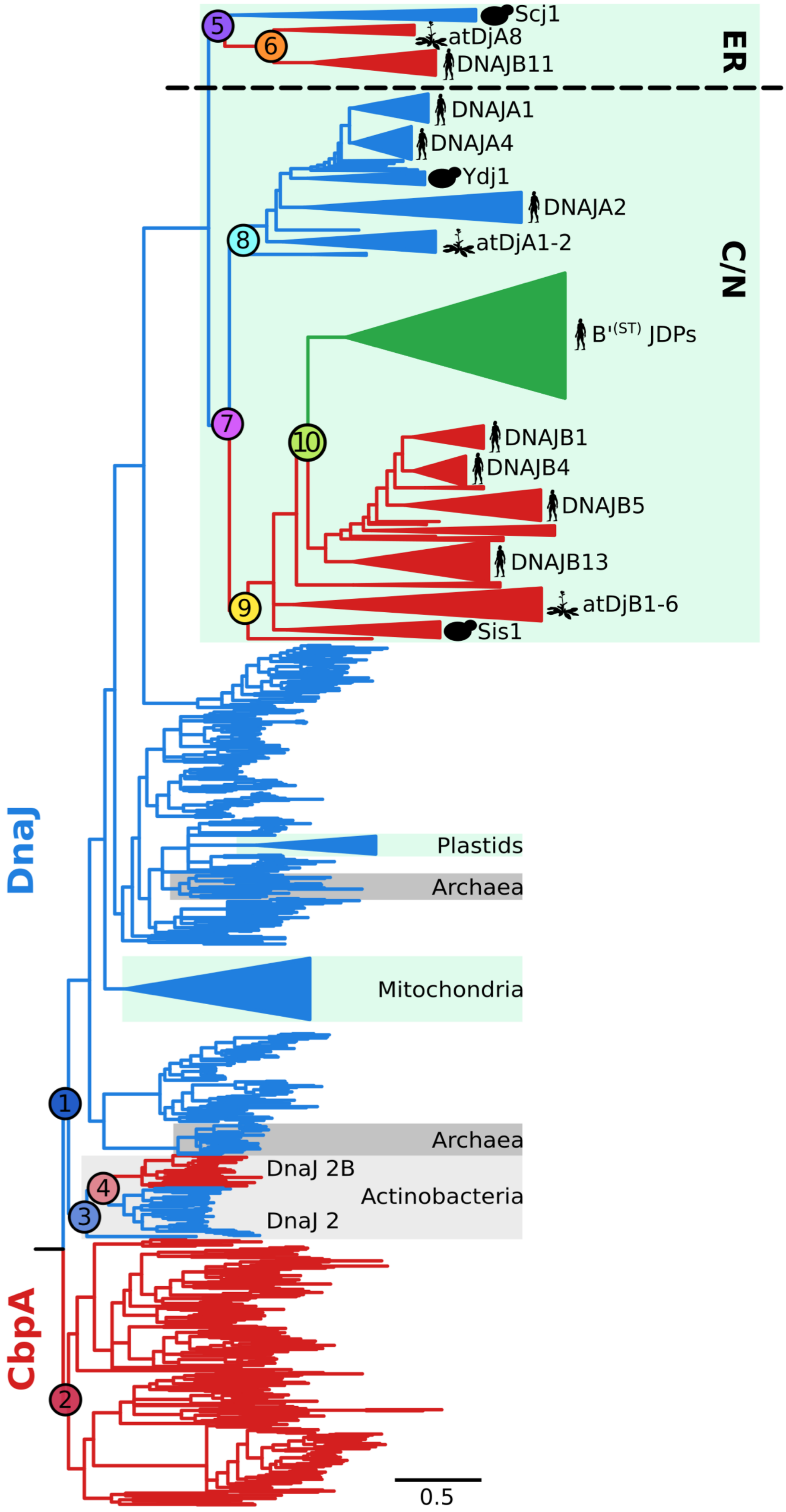


| jackknife 60 | |
|---|---|
| 1 | 98 |
| 2 | 98 |
| 3 | 99 |
| 4 | 97 |
| 5 | 100 |
| 6 | 100 |
| 7 | 99 |
| 8 | 81 |
| 9 | 98 |
| 10 | 84 |

**Figure S2.** Jackknife ML analysis of 100 replicates, each comprising 60% of randomly sampled positions from the A/B$^{C}$B'$^{(ST)}$-alignment, using LG+I+G model of sequence evolution. Color codes as in figure S1. Scale bar – amino acid substitutions per position. (right) Jackknife support values for major splits.

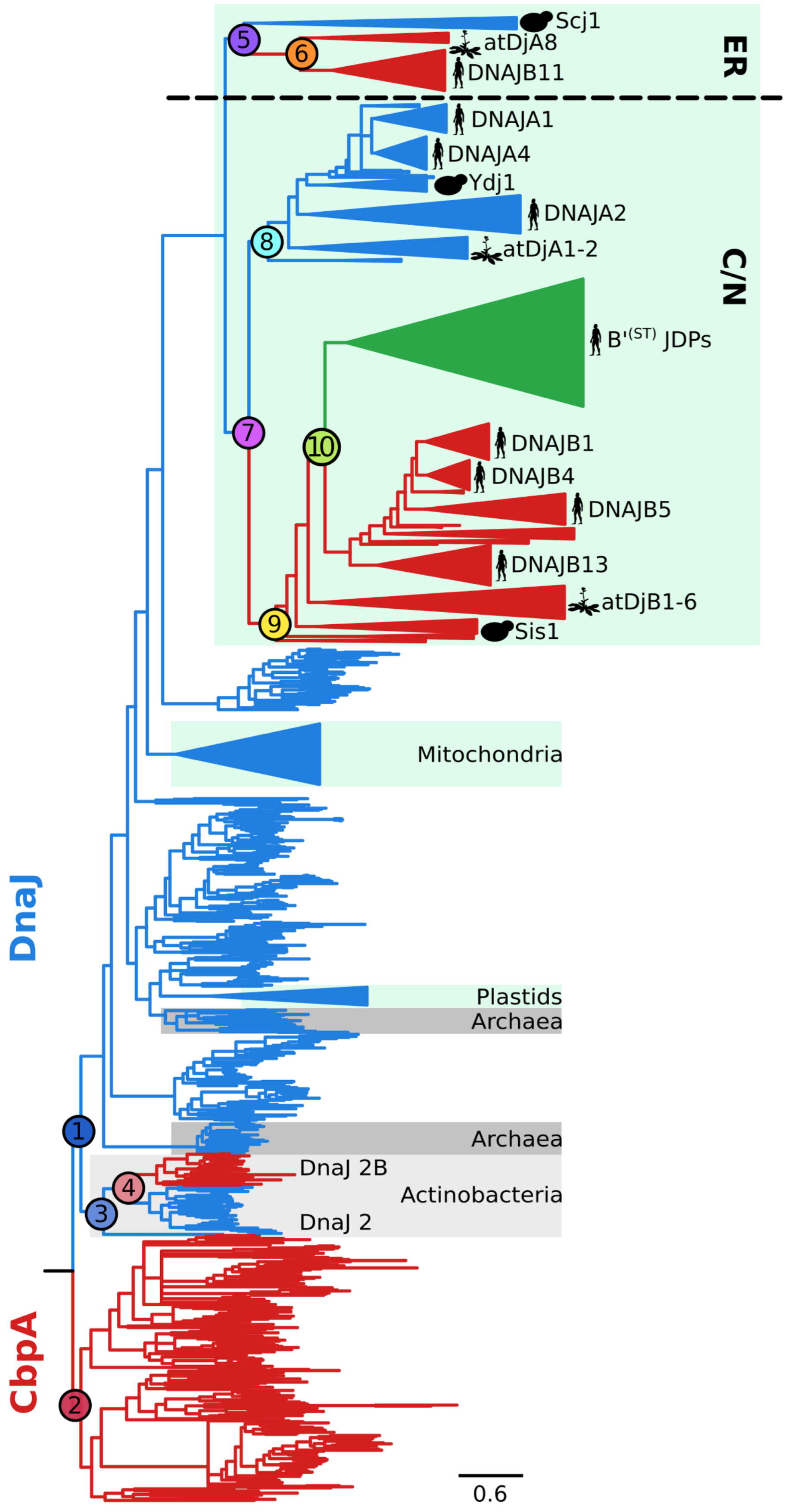


| C10+LG+G | |
|---|---|
| 1 | 92 |
| 2 | 92 |
| 3 | 100 |
| 4 | 100 |
| 5 | 100 |
| 6 | 100 |
| 7 | 100 |
| 8 | 99 |
| 9 | 100 |
| 10 | 74 |

**Figure S3.** ML phylogeny based on AB$^{C}$B'$^{(ST)}$ -alignment, reconstructed with C10 + LG +G mixture model of sequence evolution. Color codes as in figure S1. Scale bar – amino acid substitutions per position. (right) Bootstrap support values for major splits.

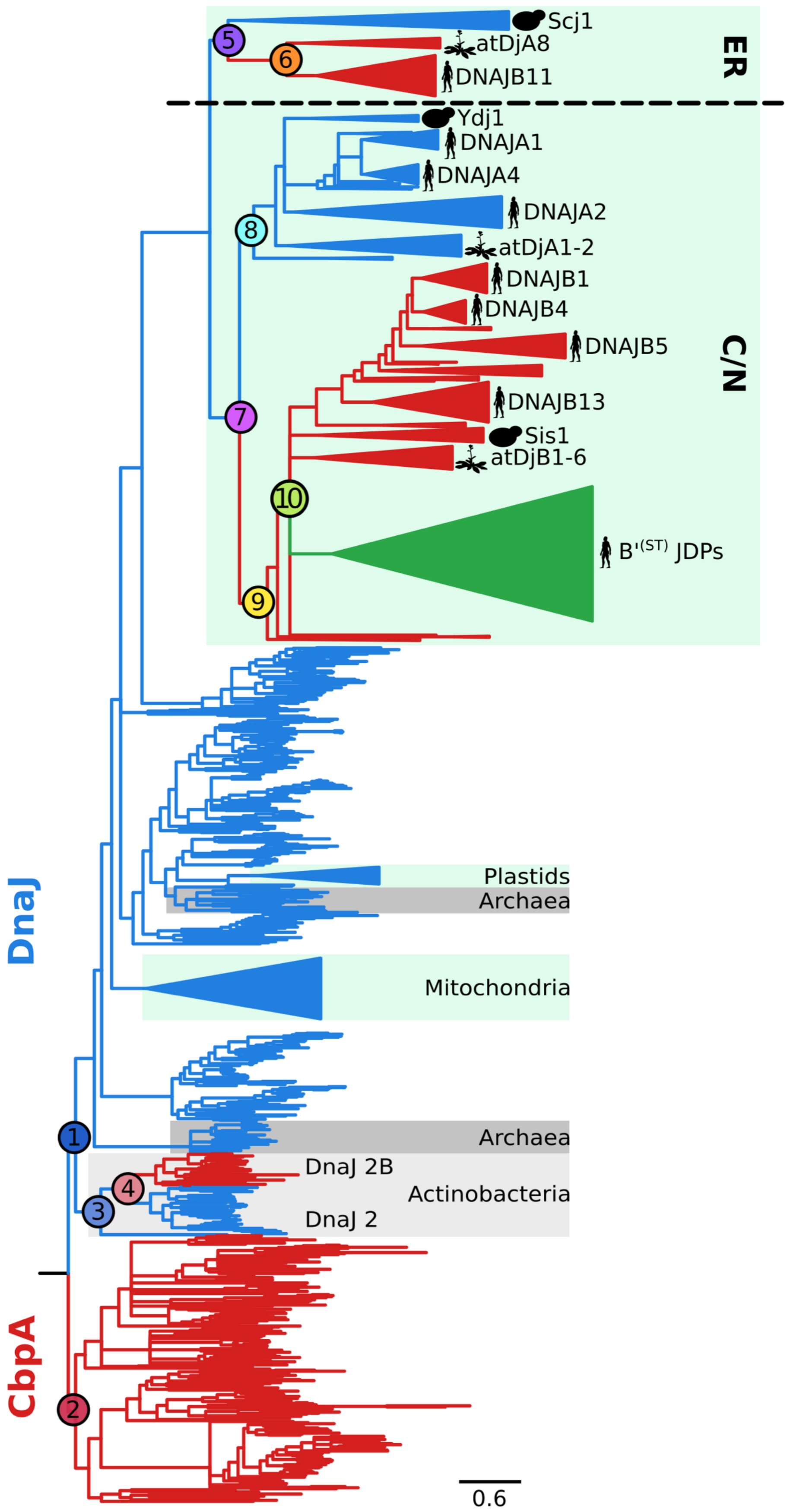


| C20+LG+G | |
|---|---|
| 1 | 94 |
| 2 | 94 |
| 3 | 100 |
| 4 | 100 |
| 5 | 100 |
| 6 | 100 |
| 7 | 100 |
| 8 | 96 |
| 9 | 100 |
| 10 | p |

**Figure S4.** ML phylogeny based on $AB^{C}B'^{(ST)}$ -alignment, reconstructed with C20 + LG +G mixture model of sequence evolution. Color codes as in figure S1. Scale bar – amino acid substitutions per position. (right) Boostrap support values for major splits. Note polytomy of the $B'^{(ST)}$ split (split 10).

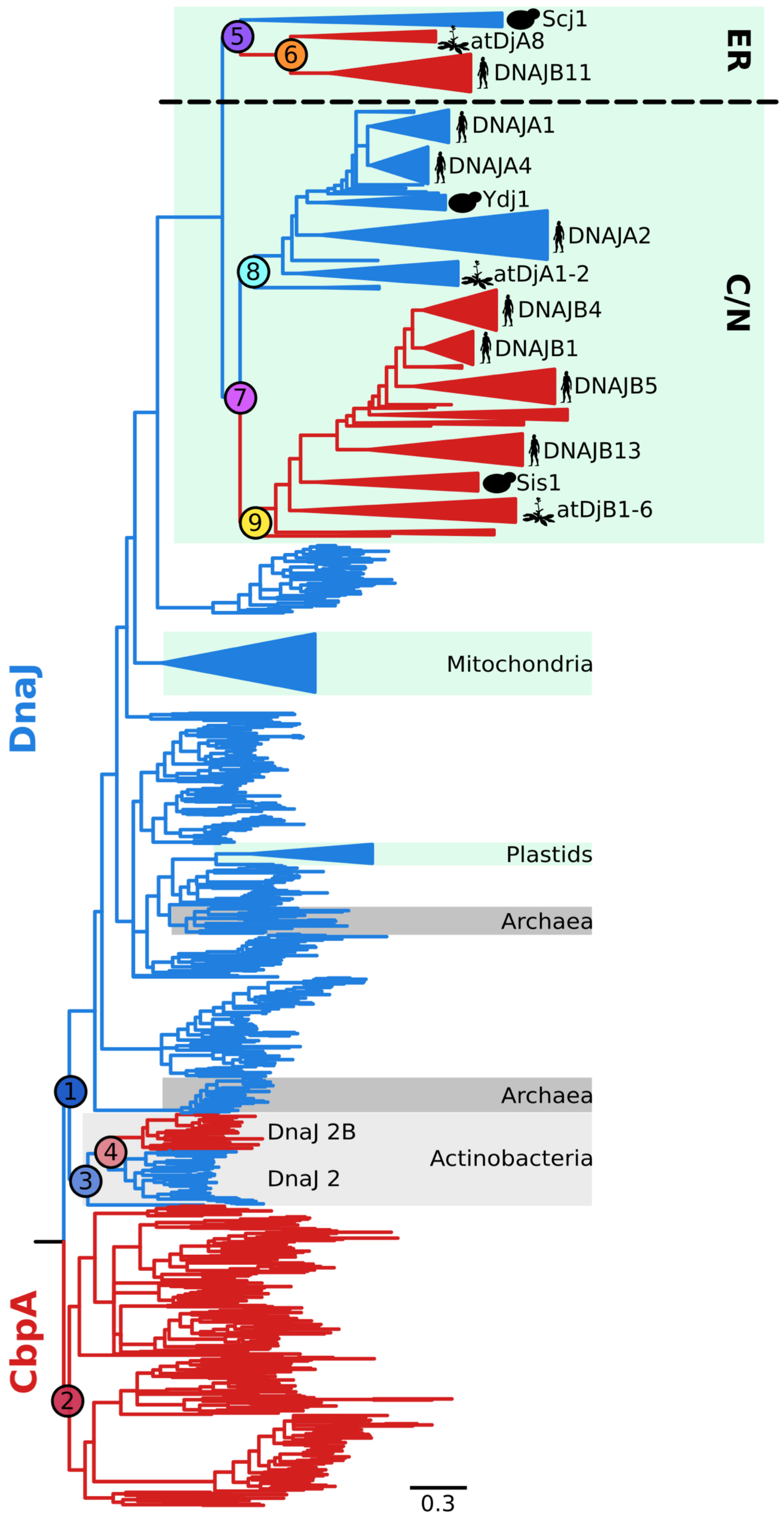


| LG+I+G | |
|---|---|
| 1 | 95 |
| 2 | 95 |
| 3 | 98 |
| 4 | 100 |
| 5 | 100 |
| 6 | 100 |
| 7 | 99 |
| 8 | 88 |
| 9 | 98 |

**Figure S5.** ML phylogeny based on the A/B$^{C}$-alignment and C20+LG+G mixture model of sequence evolution. Color codes as in figure S1. Scale bar – amino acid substitutions per position. (right) Bootstrap support for major splits.

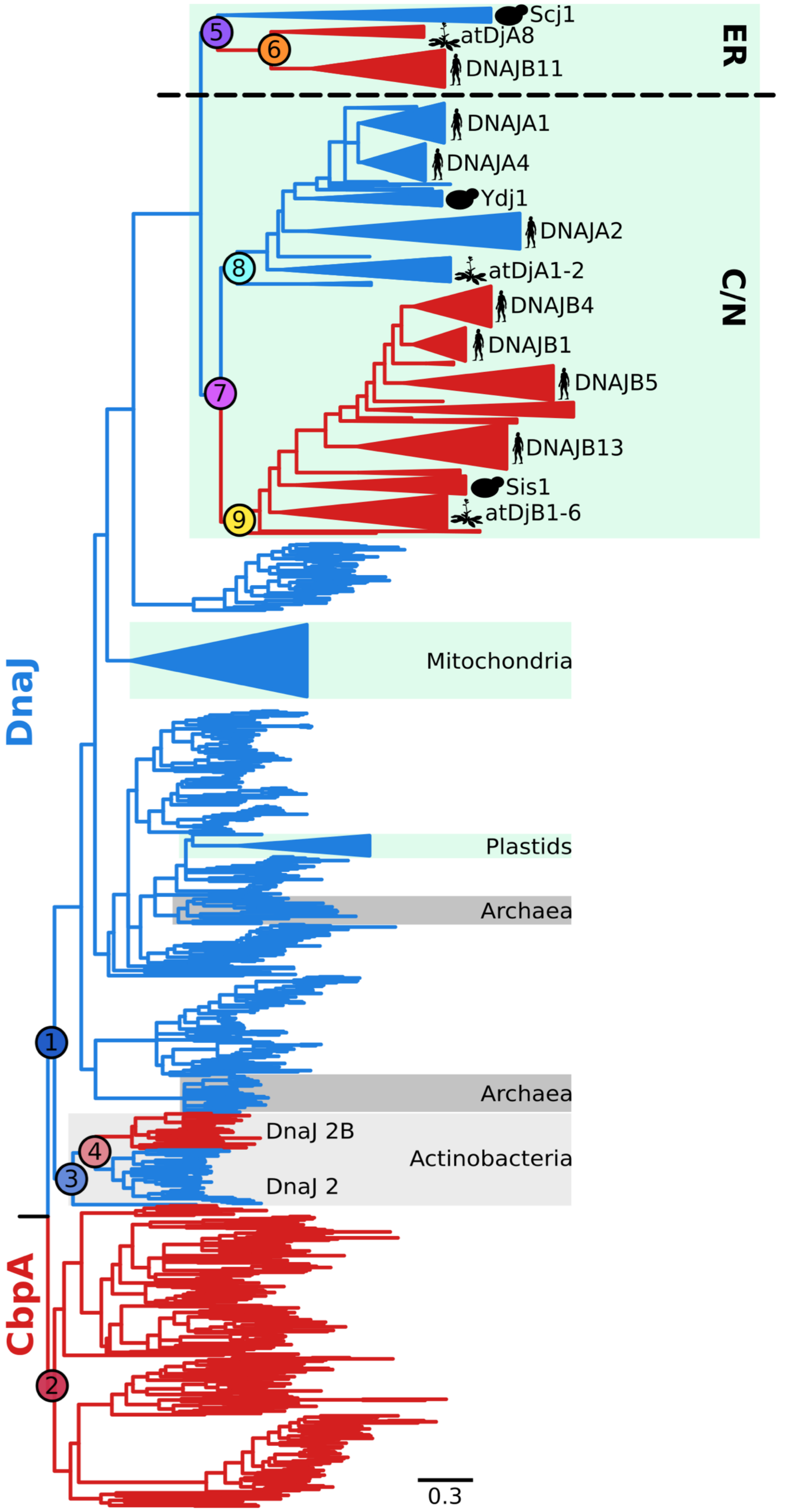


| jackknife 60 | |
|---|---|
| 1 | 80 |
| 2 | 80 |
| 3 | 99 |
| 4 | 100 |
| 5 | 100 |
| 6 | 100 |
| 7 | 99 |
| 8 | 96 |
| 9 | 100 |

**Figure S6.** ML phylogeny based on the A/B$^{C}$-alignment and C10+LG+G mixture model of sequence evolution. Color codes as in figure S1. Scale bar – amino acid substitutions per position. (right) Bootstrap support for major splits.

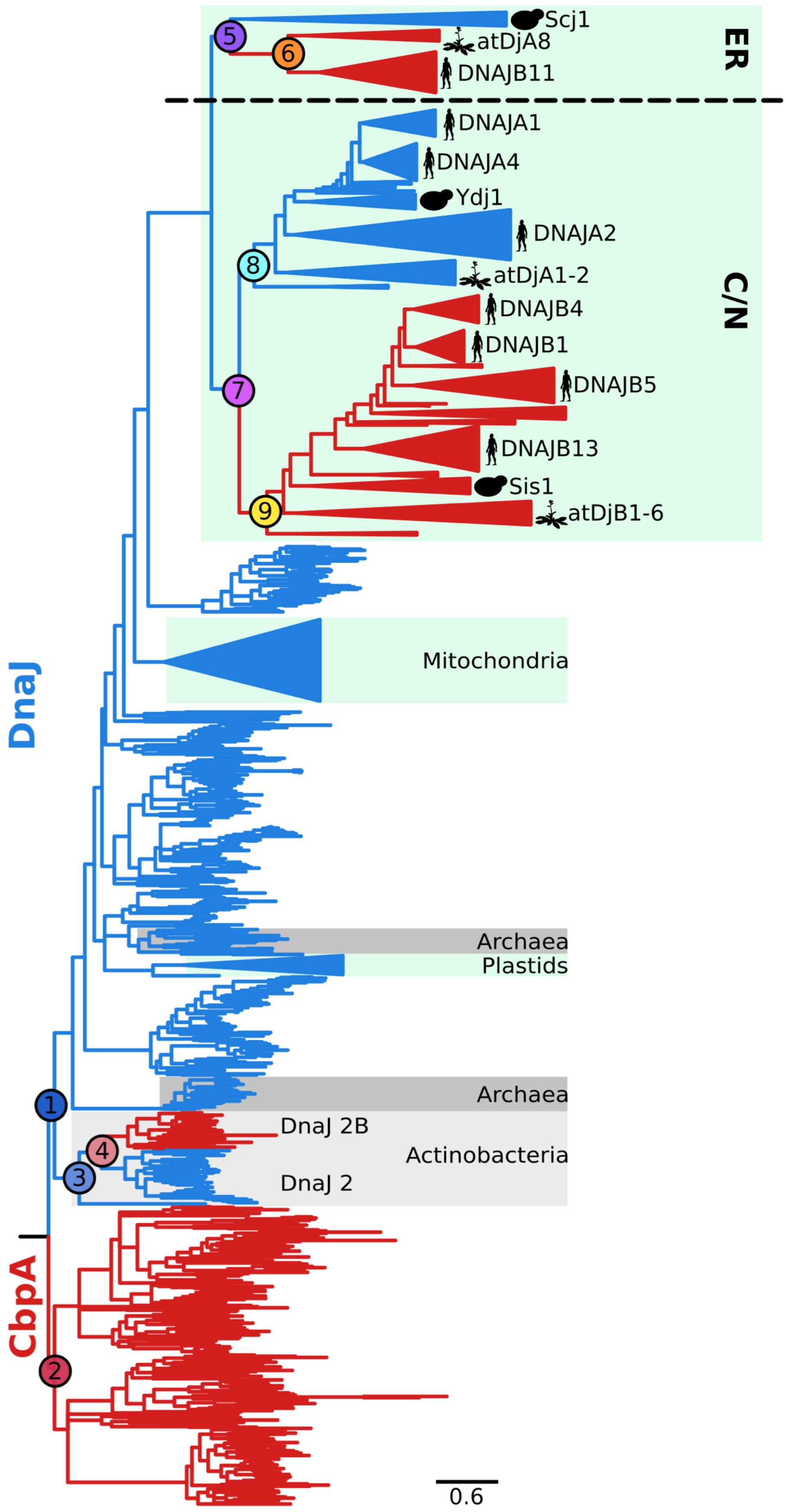


| C10+LG+G | |
|---|---|
| 1 | 88 |
| 2 | 88 |
| 3 | 99 |
| 4 | 100 |
| 5 | 100 |
| 6 | 100 |
| 7 | 100 |
| 8 | 99 |
| 9 | 100 |

**Figure S7.** Jackknife ML analysis of 100 replicates, each comprising 60% of randomly sampled positions from the A/B$^{C}$-alignment, using LG+I+G model of sequence evolution. Color codes as in figure S1. Scale bar – amino acid substitutions per position. (right) Jackknife support for major splits.

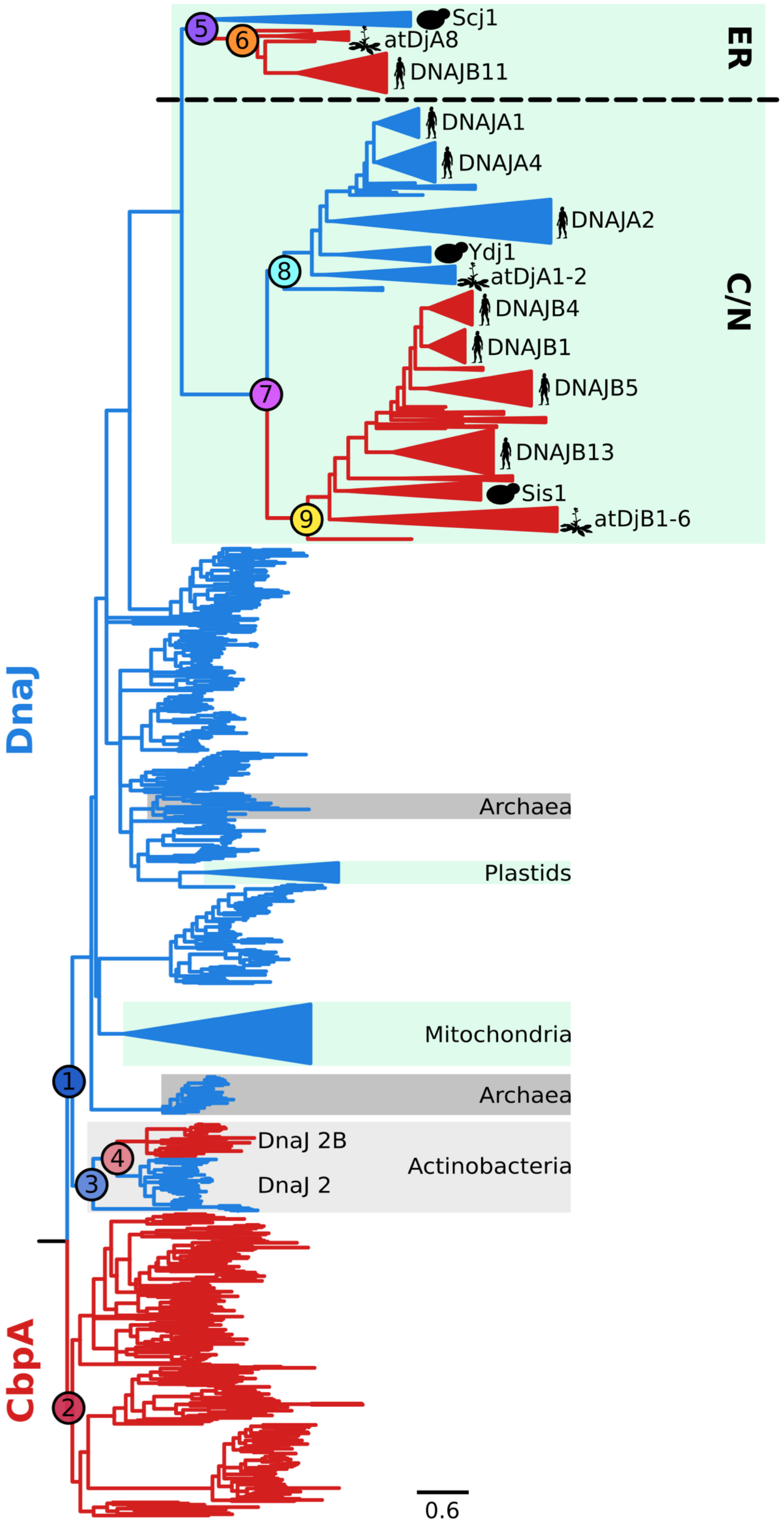


| C20+LG+G | |
|---|---|
| 1 | 89 |
| 2 | 89 |
| 3 | 100 |
| 4 | 100 |
| 5 | 100 |
| 6 | 100 |
| 7 | 100 |
| 8 | 100 |
| 9 | 100 |

**Figure S8.** ML phylogeny based on the A/B$^{C}$-alignment and LG+I+G model of sequence evolution. Color codes as in figure S1. Scale bar – amino acid substitutions per position. (right) Bootstrap support for major splits.

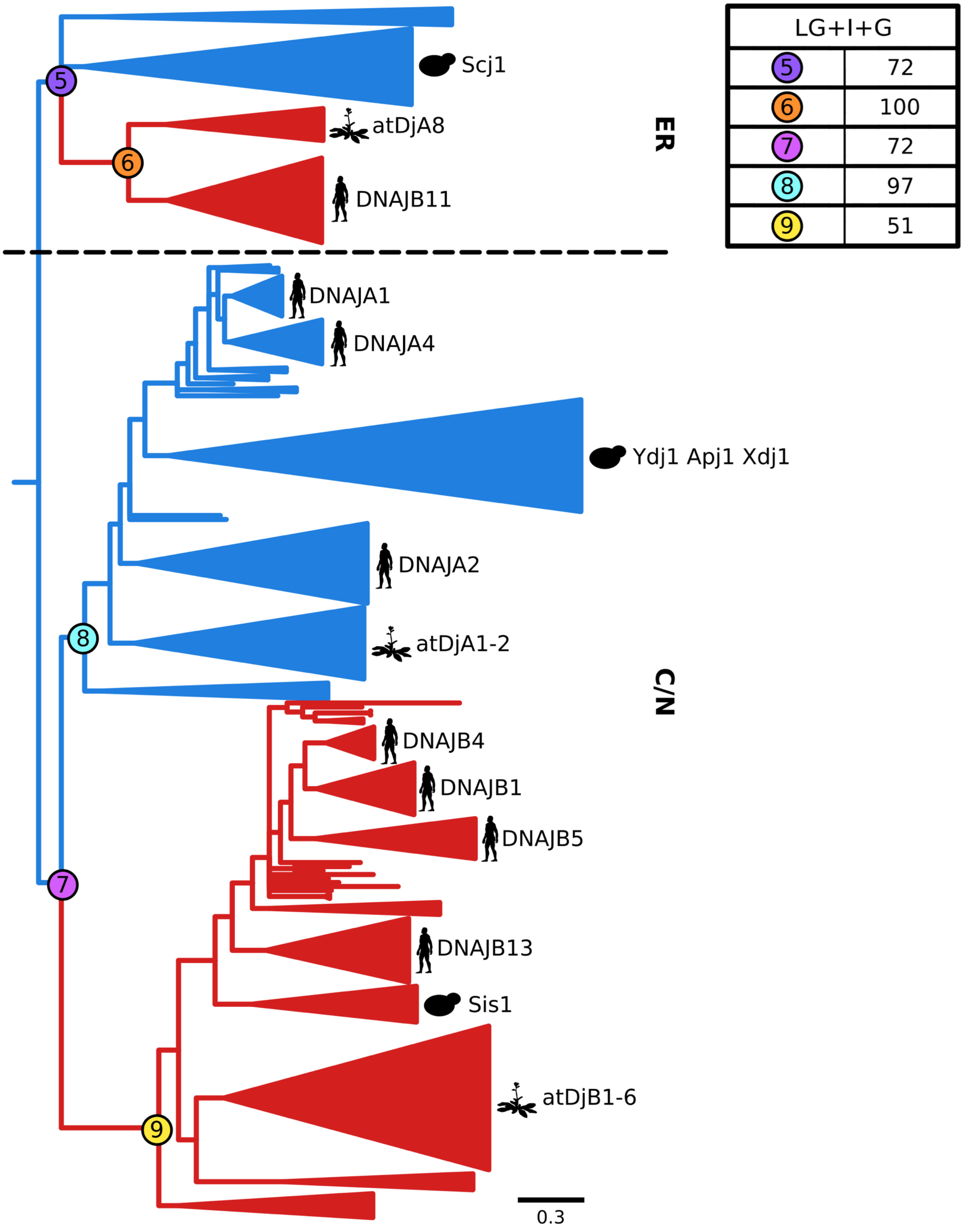


**Figure S9.** Bayesian phylogeny based on the A/B$^{C}$E alignment and WLSR5 model of sequence evolution. Color codes as in figure S1. Scale bar – amino acid substitutions per position. (right) Posterior probability for major splits.

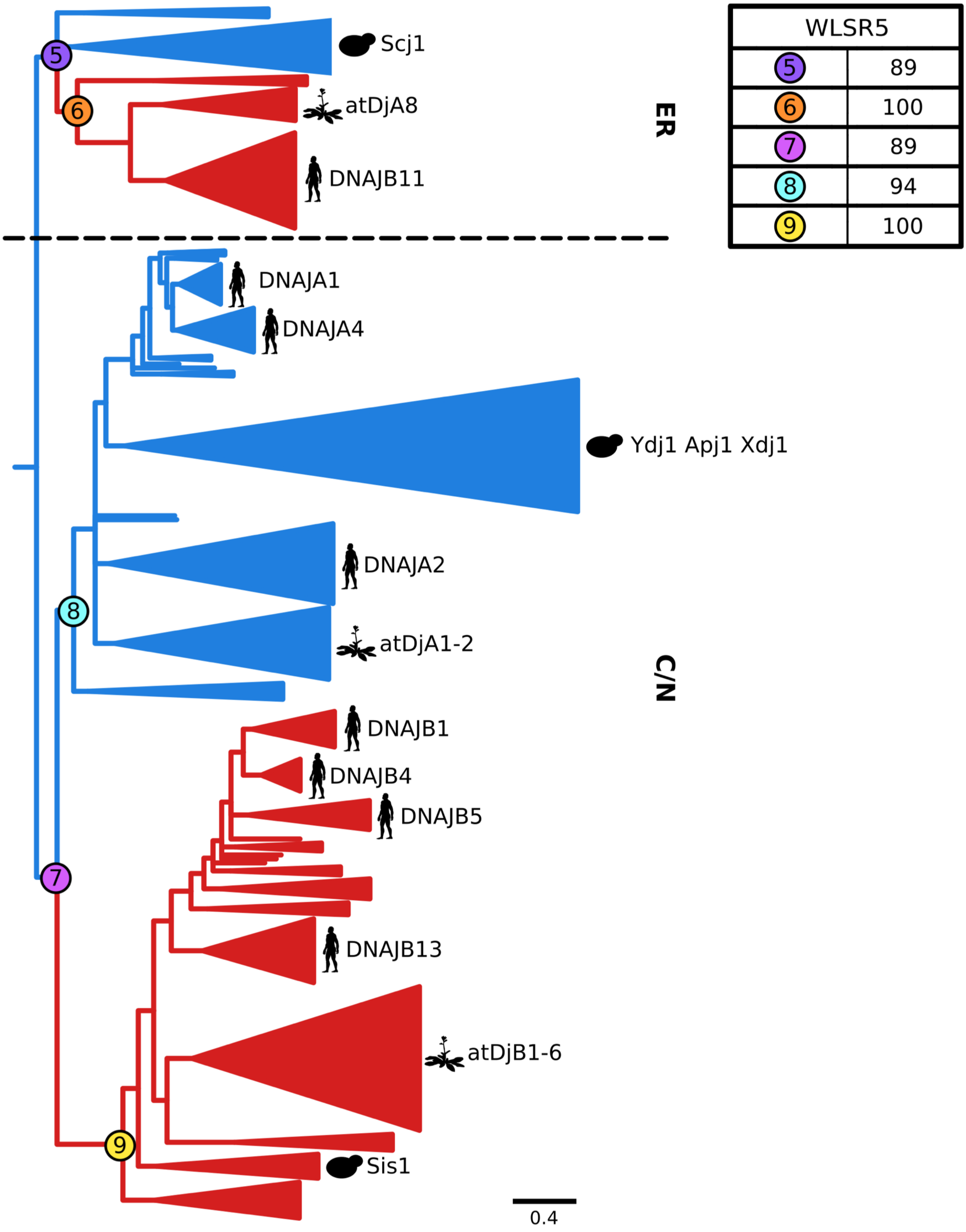


**Figure S10.** Bayesian phylogeny based on the A/B$^{C}$E alignment and LG+I+G model of sequence evolution. Color codes as in figure S1. Scale bar – amino acid substitutions per position. (right) Posterior probability for major splits.

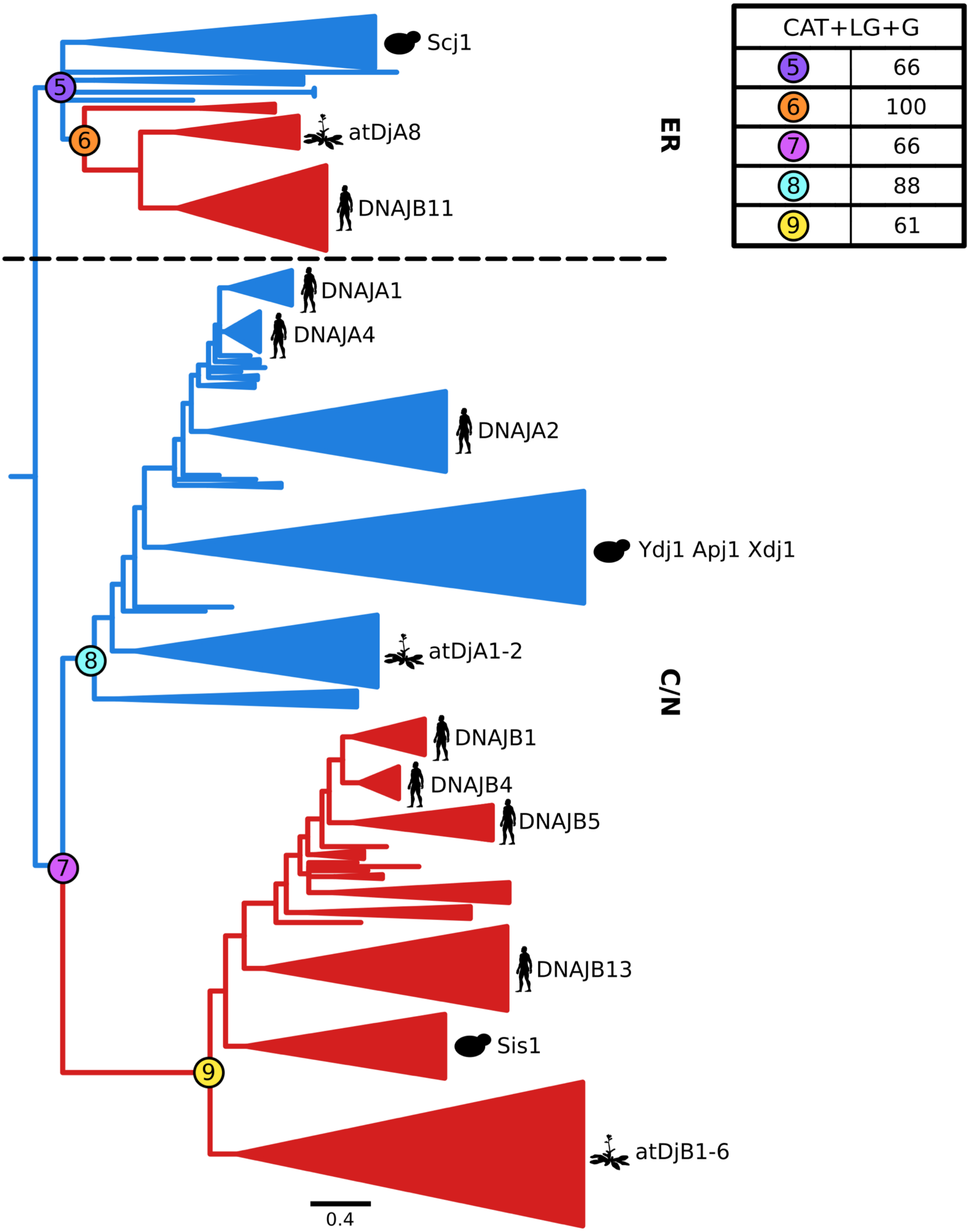


**Figure S11.** Bayesian phylogeny based on the A/B$^{C}$E alignment and C60+LG+G model of sequence evolution. Color codes as in figure S1. Scale bar – amino acid substitutions per position. (right) Posterior probability for major splits.

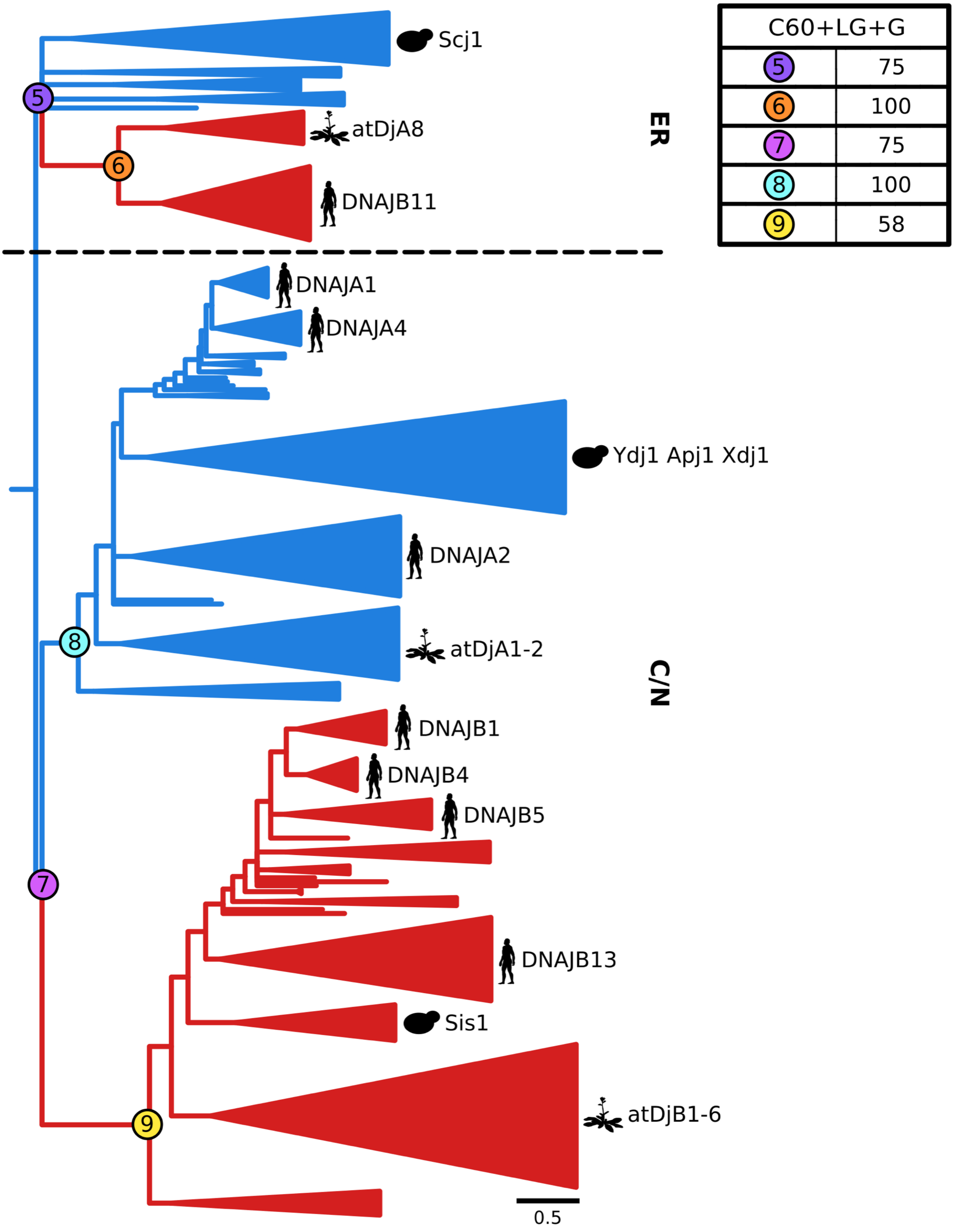


**Figure S12.** Bayesian phylogeny based on the A/B$^{C}$E alignment and CAT+LG+G model of sequence evolution. Color codes as in figure S1. Scale bar – amino acid substitutions per position. (right) Posterios probability for major splits.

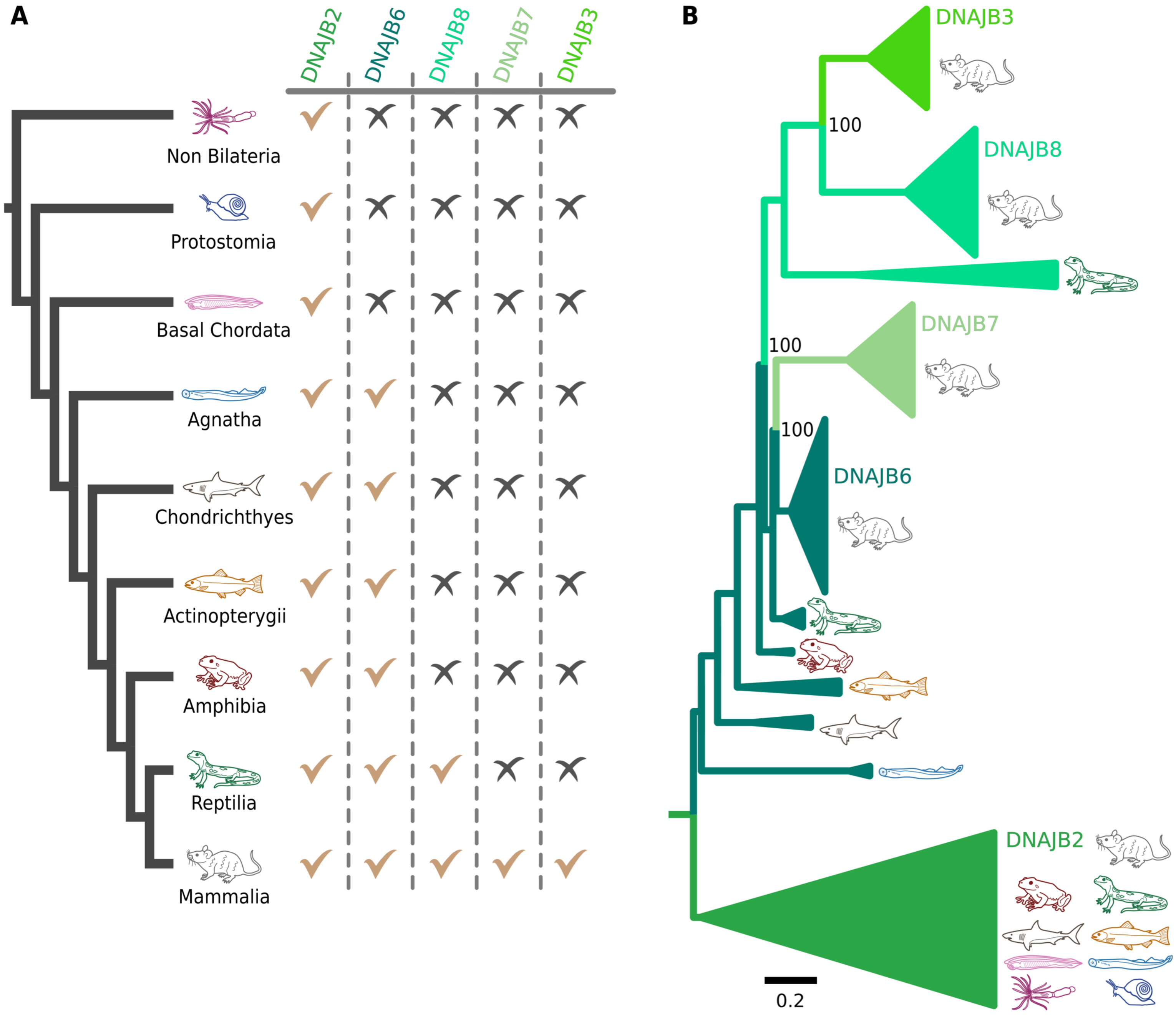


**Figure S13.** Evolution of B'(ST) from the cytosol/nucleus of metazoans. (A) Phylogenetic distribution of B'(ST) JDPs across major metazoan clades; presence ( ) or absence (✗). (B) ML phylogeny based on B'(ST) alignment, reconstructed with JTT+I+G4 model of sequence evolution. The tree was constrained to enforce monophyly of a clade encompassing DNAJB8/B6/B3 homologs from reptiles and mammals. Using the approximately unbiased (AU) test constrained topology has a higher score than unconstrained topology (0.552 vs 0.448). Different shades of green correspond to each B'(ST) paralog. Scale bar – amino acid substitutions per position.

**A**

| | J-domain A | J-domain $B^{C}$ |
|---|---|---|
| J-domain $B'^{(ST)}$ | $7.6*10^{-45}$<br>214.6 | $1.9*10^{-50}$<br>240.3 |

| | G/F region A | G/F region $B^{C}$ |
|---|---|---|
| G/F region $B'^{(ST)}$ | $6*10^{-4}$<br>19.6 | $1.3*10^{-10}$<br>47.5 |

**B**

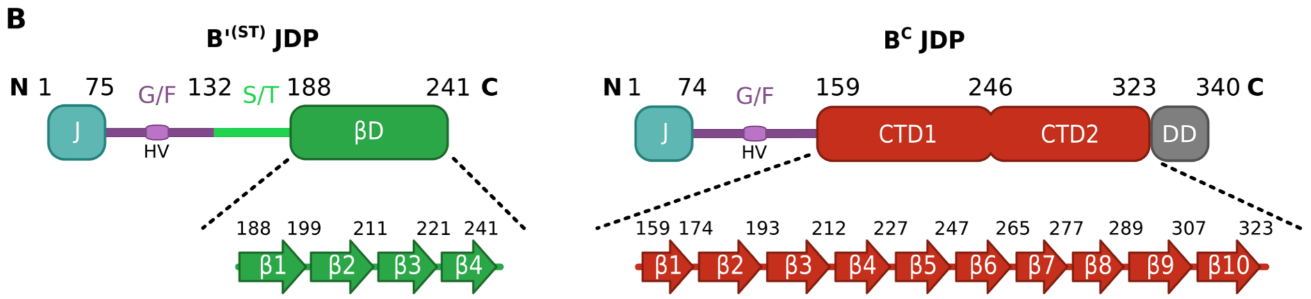


**C**

| | CTD1 $B^{C}$ | CTD2 $B^{C}$ | CTD2\|DD $B^{C}$ | CTD1\|CTD2 $B^{C}$ | CTD1\|CTD2\|DD $B^{C}$ |
|---|---|---|---|---|---|
| S/T\|βD $B'^{(ST)}$ | 0.59<br>8.8 | 0.43<br>9.5 | 0.49<br>9.4 | 0.46<br>10.8 | 0.47<br>10.9 |
| S/T $B'^{(ST)}$ | 0.93<br>5.3 | 0.76<br>6.3 | 0.78<br>6.5 | 0.92<br>6.3 | 0.92<br>6.4 |
| βD $B'^{(ST)}$ | 0.3<br>9.3 | 0.11<br>11.2 | 0.12<br>11.3 | 0.25<br>11.0 | 0.26<br>11.1 |

**D**

| | β1 $B^{C}$ | β2 $B^{C}$ | β3 $B^{C}$ | β4 $B^{C}$ | β5 $B^{C}$ | β6 $B^{C}$ | β7 $B^{C}$ | β8 $B^{C}$ | β9 $B^{C}$ | β10 $B^{C}$ |
|---|---|---|---|---|---|---|---|---|---|---|
| β1 $B'^{(ST)}$ | 0.31<br>3.0 | 0.18<br>5.1 | 0.35<br>5.1 | 0.20<br>4.3 | 0.55<br>3.0 | 0.55<br>2.8 | 0.20<br>2.6 | 0.073<br>3.1 | 0.25<br>4.2 | 0.41<br>3.5 |
| β2 $B'^{(ST)}$ | 0.22<br>3.5 | 0.037<br>7.0 | 0.21<br>5.9 | 0.023<br>6.8 | 0.30<br>3.9 | 0.066<br>5.3 | 0.097<br>3.4 | 0.051<br>3.5 | 0.13<br>5.0 | 0.28<br>4.1 |
| β3 $B'^{(ST)}$ | 0.34<br>2.8 | 0.91<br>2.3 | 0.54<br>4.3 | 0.84<br>2.1 | 0.37<br>3.5 | 0.62<br>2.5 | 0.11<br>3.1 | 0.31<br>1.6 | 0.12<br>4.8 | 0.82<br>2.2 |
| β4 $B'^{(ST)}$ | 0.32<br>2.6 | 0.13<br>5.0 | 0.23<br>5.2 | 0.64<br>2.5 | 0.32<br>3.3 | 0.55<br>2.4 | 0.039<br>3.7 | 0.023<br>3.7 | 0.20<br>4.0 | 0.098<br>4.7 |

**Figure S14.** Profile-profile comparisons between $B'^{(ST)}$ and class A and $B^{C}$ JDPs.
(A) Profile-profile comparisons of J-domain and G/F between $B'^{(ST)}$ and A or $B^{C}$ JDPs - E-values (top) and HHalign scores (bottom). (B) Schematic representation of domain organization in $B'^{(ST)}$ and $B^{C}$ JDPs, with highlighted regions used in the profile-profile comparisons shown in (C) and (D). (C) Profile-profile comparisons between S/T and βD of $B'^{(ST)}$s and CTD1/2 of $B^{C}$s, and (D) between β-strands from βD of $B'^{(ST)}$ and from CTD1 and CTD2 domains of $B^{C}$s - E-values (top) and HHalign scores (bottom).

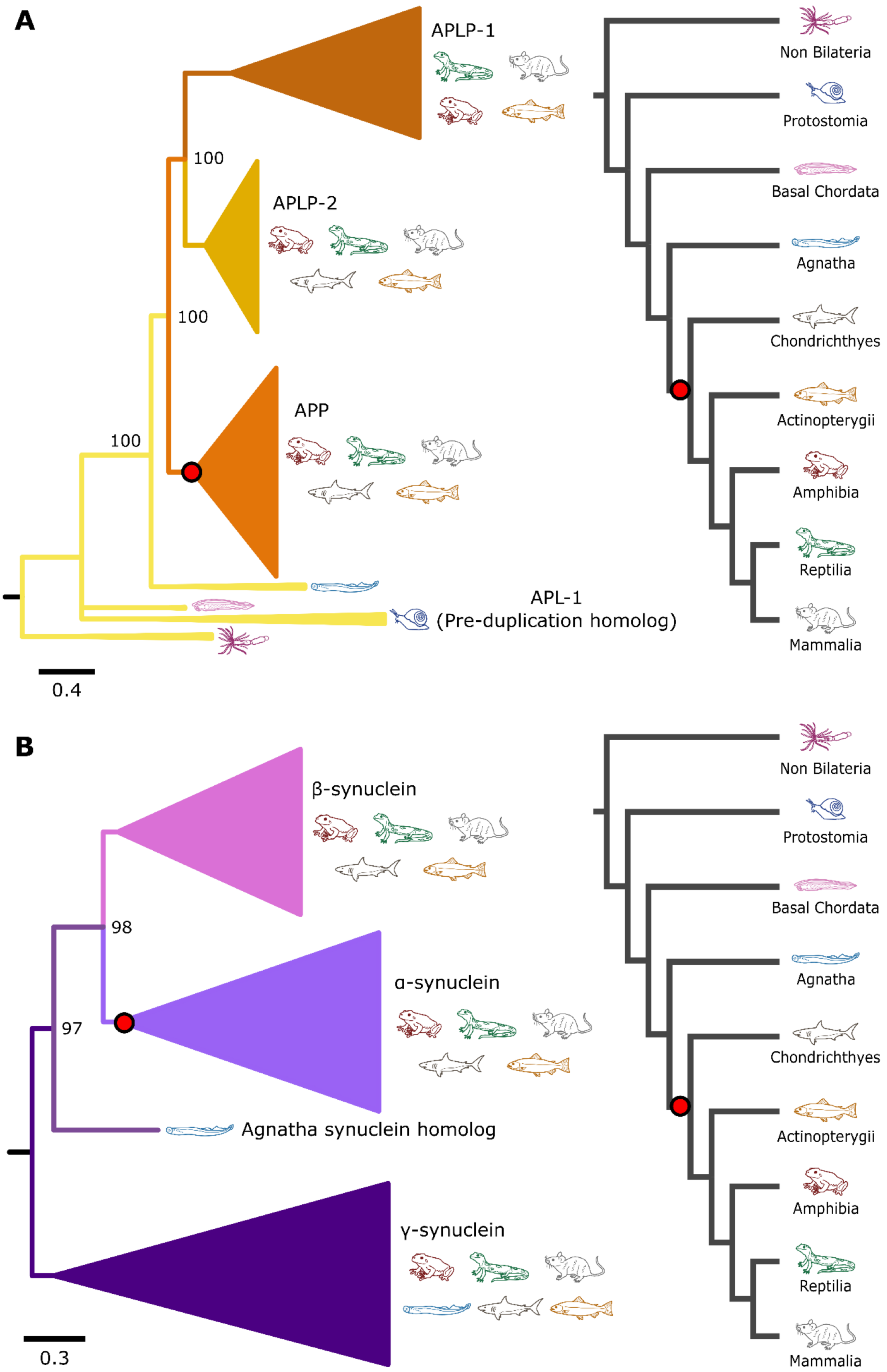


**Figure S15.** Phylogenetic analysis of APP and synuclein homologs from metazoans. Protein trees (left) and simplified phylogeny of metazoans (right). Inferred emergence of precursor of human amyloidogenic proteins indicated by red dots. (A) ML phylogeny of 545 APP homologs: APL-1, APLP-1, APLP-2 and APP reconstructed using the JTT+I+G4 model. (B) ML phylogeny of 348 synuclein homologs: α- β- and γ- synuclein and Agnatha synuclein homolog, reconstructed using the JTT+G4 model. Bootstrap support for major splits is indicated. Splits with support <50 were collapsed into polytomies. Scale bars – amino acid substitutions per position.

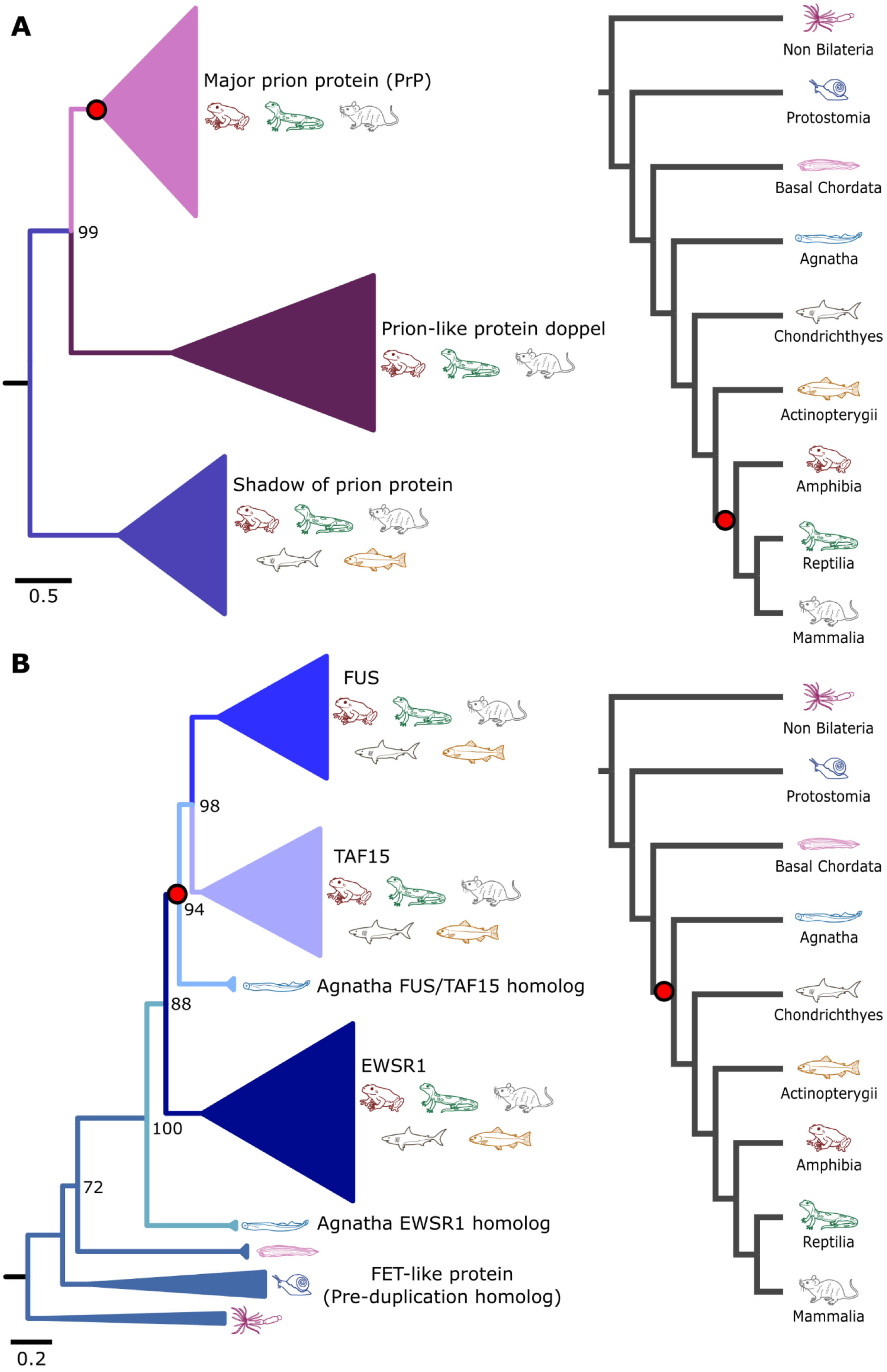


**Figure S16.** Phylogenetic analysis of major prion protein (PrP) and FUS/TAF15 homologs from metazoans. Protein trees (left) and simplified phylogeny of metazoans (right). Inferred emergence of precursor of human amyloidogenic proteins indicated by red dots. (A) ML phylogeny of 209 major prion protein homologs: PrP, prion like protein doppel, shadow of prion protein reconstructed using the JTT+F+G4 model. (B) ML phylogeny of 426 FUS/TAF15 homologs: FUS, TAF15, EWSR1 and FET like protein reconstructed using the JTT+F+G4 model. Bootstrap support for major splits is indicated. Splits with support <50 were collapsed into polytomies. Scale bars – amino acid substitutions per position.

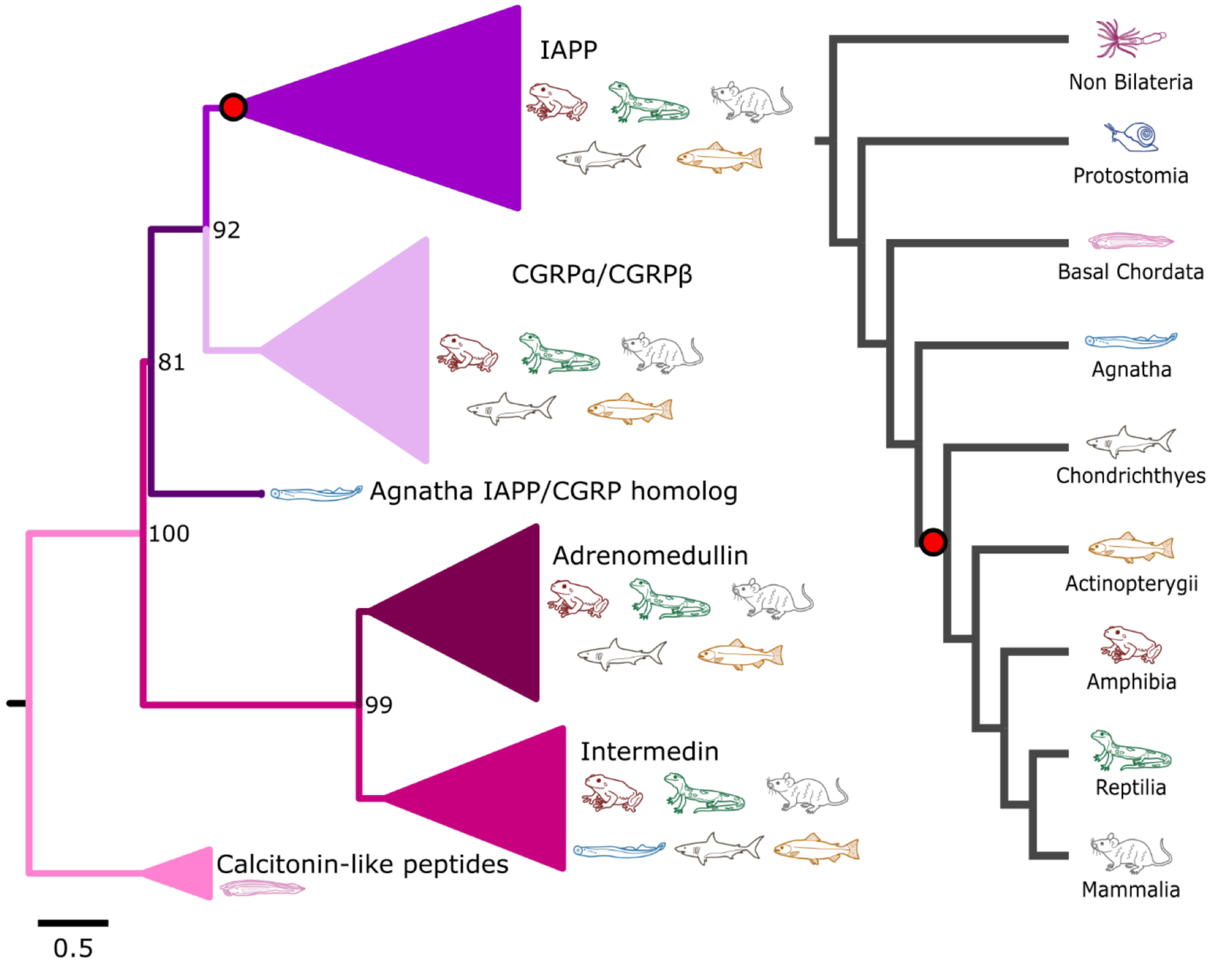


**Figure S17.** Phylogenetic analysis of 514 IAPP homologs: IAPP, CGRP$\alpha$ and $\beta$, adrenomedullin, intermedin, calcitonine like peptides. Protein tree (left) and simplified phylogeny of metazoans (right). Inferred emergence of precursor of human amyloidogenic IAPP indicated by red dot. The ML tree was reconstructed using JTT+G4 model. Bootstrap support for major splits is indicated. Splits with support <50 were collapsed into polytomies. Scale bar – amino acid substitutions per position.

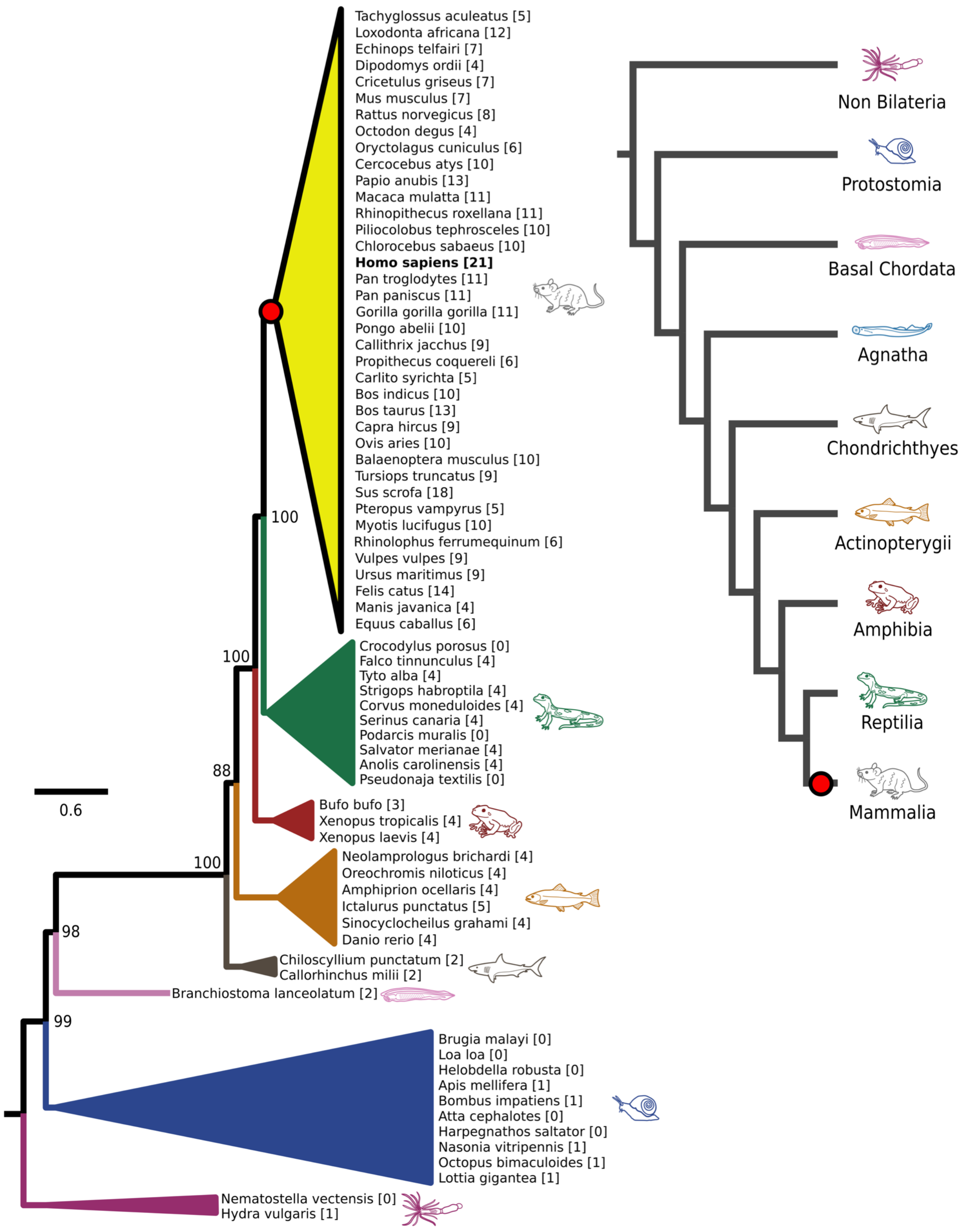


**Figure S18.** Phylogenetic analysis of 59 huntingtin protein (HTT) orthologs from metazoans. Protein tree (left) simplified metazoans phylogeny (right). Inferred emergence of HTT-polyQ indicated by red dot. The length of polyQ motifs indicated in square brackets. Bootstrap support for major splits is indicated. Nodes with support <50 were collapsed into polytomies. Scale bar – amino acid substitutions per position.

```
1       JD                                                                       80
DNAJA2  MANVADTKLYDILGVPPGASENELKKAYRKLAKEYHPDKNP---NAGDKFKEISFAYEVLSNPEKRELYDRYGEQGLREG
AncA    --MVKETDYYDILGVSPNASEDEIKKAYRKLALKYHPDKNPDNPEAEEKFKEISEAYEVLSDPEKREIYDQYGEEGLKEG
AncAB   ----MAKDYYDILGVSRGASEDEIKKAYRKLAMKYHPDKNPD-PEAEEKFKEISEAYEVLSDPEKREIYDQYGEEGLKGG
AncB    ----MGKDYYDILGVSRNASEDEIKKAYRKLAMKYHPDKNPD-PEAEEKFKEISEAYEVLSDPEKREIYDQYGEEGLKGG
DNAJB1  ----MGKDYYQTLGLARGASDEEIKRAYRRQALRYHPDKNKE-PGAEEKFKEIAEAYDVLSDPRKREIFDRYGEEGLKGS

81      G/F                                                                      160
DNAJA2  SGGGG--------------------------------------GMDDIFSHIFGGGLFG---------------------
AncA    GGG----GGGGGGGGGFPM--DDFFPGGGGGGG--------GHDAHDIFSQFFGGGGGGGGGS----GFSG---FGGGSD
AncAB   GGG-----GGGGGGGGFPG--DGFFPGGGGGGG--------SHDAHDIFSQFFGGGGGGGGGG----GFSG---FGGGGD
AncB    APPGGGGGGGGPGGSGFPGGFGGFHGGGGGGGGGGGGHGFSSRDPNDIFSQFFGGGGAGGGGFGGSSGFGFGGGGGGGDD
DNAJB1  GPSGGSGGGANGT--SFSYTF--------------------HGDPHAMFAEFFGGRNPF--------------------D

161     G/F                               CTD1                                   240
DNAJA2  ----------------------------------FMGNQSRSRNGRRRGEDMMHPLKVSLEDLYNGKTTKLQLSKNVLCS
AncA    GFFGMRGGGMGGGGGGMGGGG-----------------SRGRPRGPRKGEDIEVPLKVTLEDLYNGKTKKLKITRNVICS
AncAB   GFFGMRGGGMGGGGGGMGGGSGFSSGFRSHSG--GFAGNGRSRQPRKGPDIEVPLEVTLEDLYNGTTKKLKITRNVICS
AncB    GFFGMRGGGMGGGMGGMGGGGSGFSSGFRSHSGGFGNGSGRSGRQPRKPPPIEVPLPVTLEELYNGTTKKMKITRKRLNA
DNAJB1  TFFGQRNGEEGMDIDDPFSGFPMGMGGFTNVNF-GRSRSAQEPARKKQDPPVTHDLRVSLEEIYSGCTKKMKISHKRLN-

        ZnF                                                           CTD1       320
DNAJA2  ACSGQGGKSG-AVQKCSACRGRGVRIMIRQLAPGMVQQMQSVCSDCNGEGEVINEKDRCKKCEGKKVIKEVKILEVHVDK
AncA    KCNGKGSKGG-AVQTCSTCNGQGMKVVTRQMGPGMVQQMQQTCPKCNGKGEIINEKDRCNACHGKKVVKEKKILEVHIEP
AncAB   KCNGTGAKGG-AVQTCSTCNGQGMKVVTQQMGPGMFQQMQQTCPKCNGKGKII--KDRCKECHGKKVVKEKKILTVNIEP
AncB    S---------------------------------------------------------------GKTVKEDKILTINVKP
DNAJB1  -------------------------------------------------------------PDGKSIRNEDKILTIEVKK

        CTD1                                CTD2                                 400
DNAJA2  GMKHGQRITFTGEADQAPGVEPGDIVLLLQEKEHEVFQRDGNDLHMTYKIGLVEALCGFQFTFKHLDGRQIVVKYPPGKV
AncA    GMKDGQKITFPGEGDQ-PGIIPGDIVFVIQEKEHPRFKRKGNDLFMEHTISLTEALCGFTFTIKHLDGRTLTISSDPGDV
AncAB   GMKDGQKITFPGEGDQ-PGGIPGDIVFVIREKPHPRFKRKGNDLFYEVTISLTEALCGFTVTITHLDGRTLKISSDPGDV
AncB    GWKEGTKITFPGEGDQQPGRIPADIVFVIKEKPHPRFKRDGNDLIYTVKISLTEALCGFTVQITTLDGRTLKIPVN--DV
DNAJB1  GWKEGTKITFPKEGDQTSNNIPADIVFVLKDKPHNIFKRDGSDVIYPARISLREALCGCTVNVPTLDGRTIPVVFK--DV

        CTD2                                  DD                CTE              480
DNAJA2  IEPGCV-RVVRGEGMPQYRNPFEKGDLYIKFDVQFPENNWINPDKLSELEDLLPSRPEVPNIIGE-TEE--VE-LQEFDS
AncA    IQPGSV-KCIPGEGMPIHRNPHERGDLYIQFNVEFPNSLT-P-EQLKALKEVLPSSSSPTVPKDDKVEECELE-VDPNQY
AncAB   IQPGSV-KTIPGEGMPIHRNPHQRGDLYIQFNVEFPKSLT-P-EQKKALKEVLPS-------------------------
AncB    IQPGSEKTVIPGEGMPISKNPGQRGDLIIEFNVKFPKSLT-P-EQKKALKEVLPS-------------------------
DNAJB1  IRPGMR-RKVPGEGLPLPKTPEKRGDLIIEFEVIFPERIP--QTSRTVLEQVLPI-------------------------

        CTE                                514
DNAJA2  TRGSGGGQRREAYNDSSDEESSSHHGPGVQCAHQ
AncA    RRSSQQHASREAYDEDDDE--DG--GPGVQCAQQ
AncAB   ----------------------------------
AncB    ----------------------------------
DNAJB1  ----------------------------------
```

**Figure S19.** Sequence alignment of reconstructed ancestral JDPs. J – J-domain, teal; G/F – glycine/phenylalanine rich region, violet; CTD1/CTD2, - substrate binding domains, blue; ZnF, zinc finger, yellow; DD – dimerization domain, grey, CTE – C-terminal extension, pink.

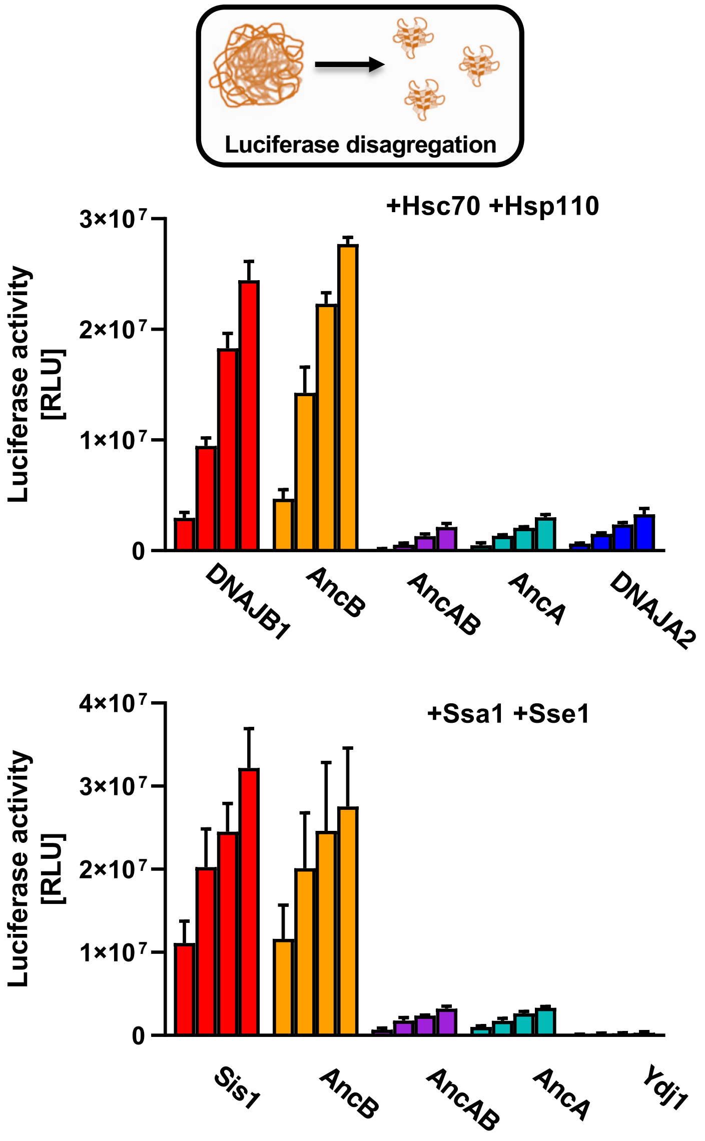


**Figure S20.** Disaggregation of luciferase aggregates (200 nM) in the presence of chaperones (top) Hsc70, 1.5 µM; Hsp105, 0.15 µM and JDP, 1 µM - as indicated.
(bottom) Ssa, 1 µM; Sse1, 0.1 µM and JDP, 1 µM - as indicated. Activity was measured after 1, 2, 3 and 4 hours. Error bars show SD from three independent repeats.

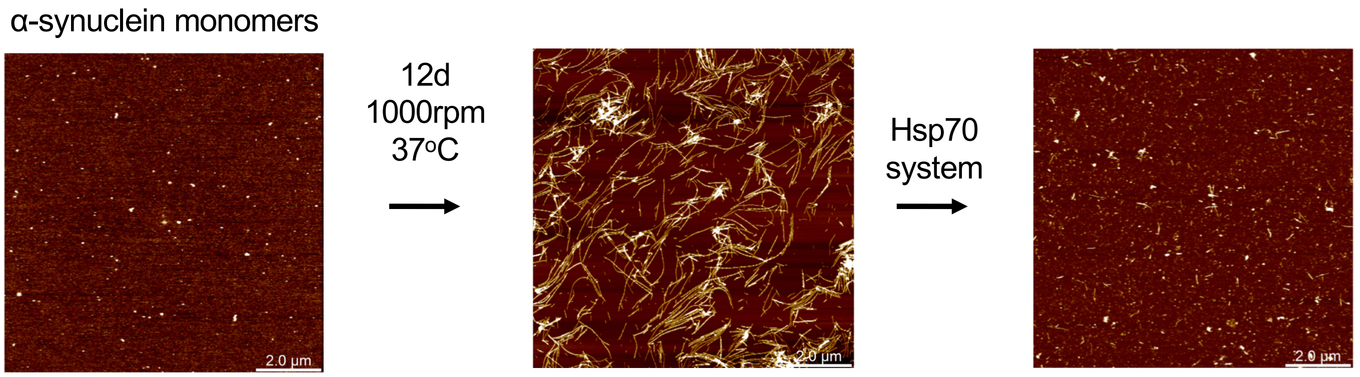


**Figure S21.** AFM images of α-synuclein recorded (left) prior to fibril formation (center) after fibril formation (12 days at 37°C with shaking) and (right) after disaggregation with Hsc70, 3 µM: DNAJB1, 0.25 µM and Hsp105, 0.3 µM.

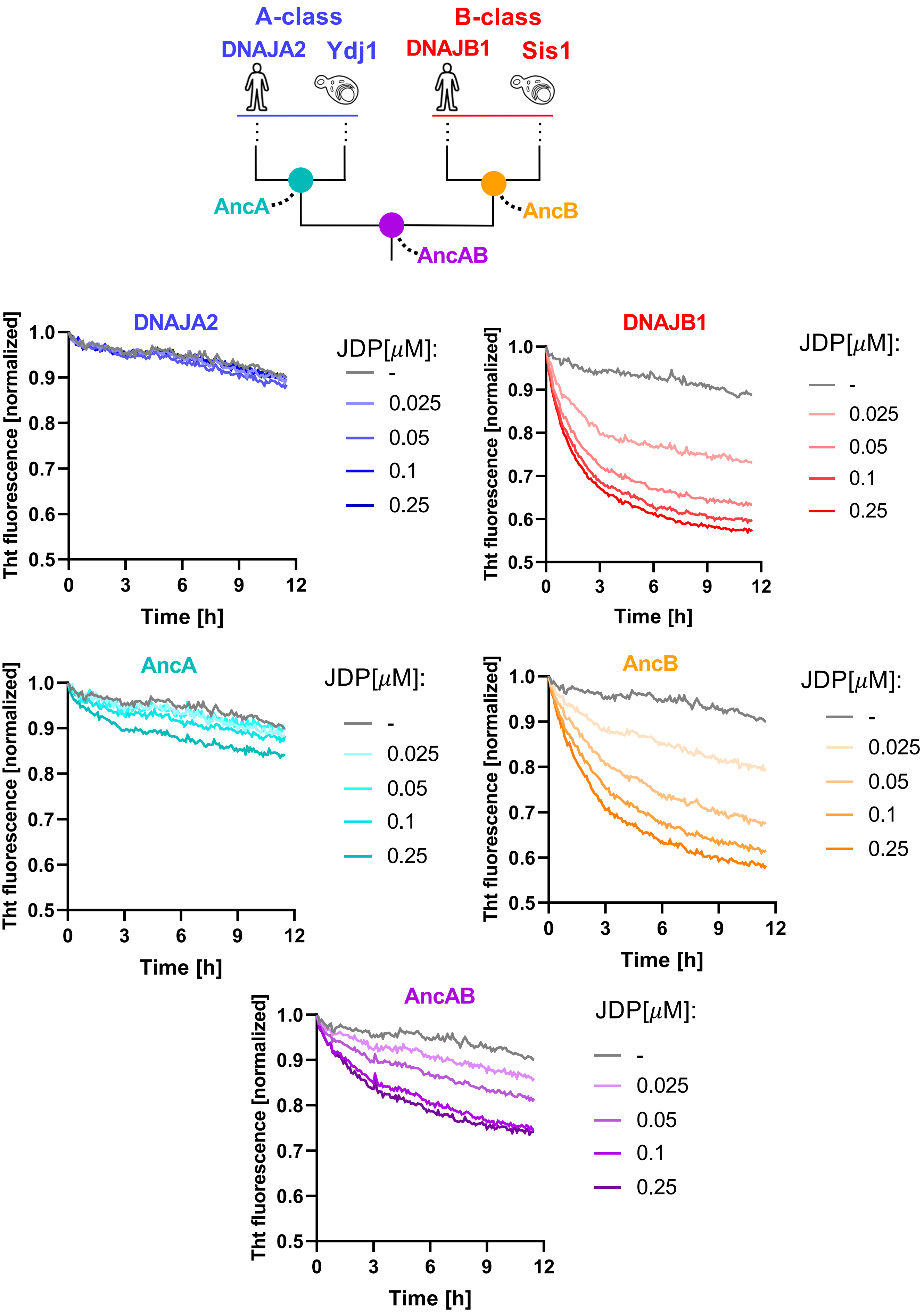


**Figure S22.** Disaggregation of α-synuclein fibrils by ancestral and contemporary JDPs.
(A) Simplified phylogeny of class A and B$^{C}$ JDPs. (B) Disassembly of α-synuclein fibrils monitored by ThT fluorescence. Reaction mixtures contained α-synuclein fibrils 0.8 μM, Hsc70 3 μM, Hsp105 0.3 μM and the indicated concentrations of the contemporary and ancestral JDPs. Data from the control experiment without JDP are the same in each graph. Traces represent mean value from three independent experiments.

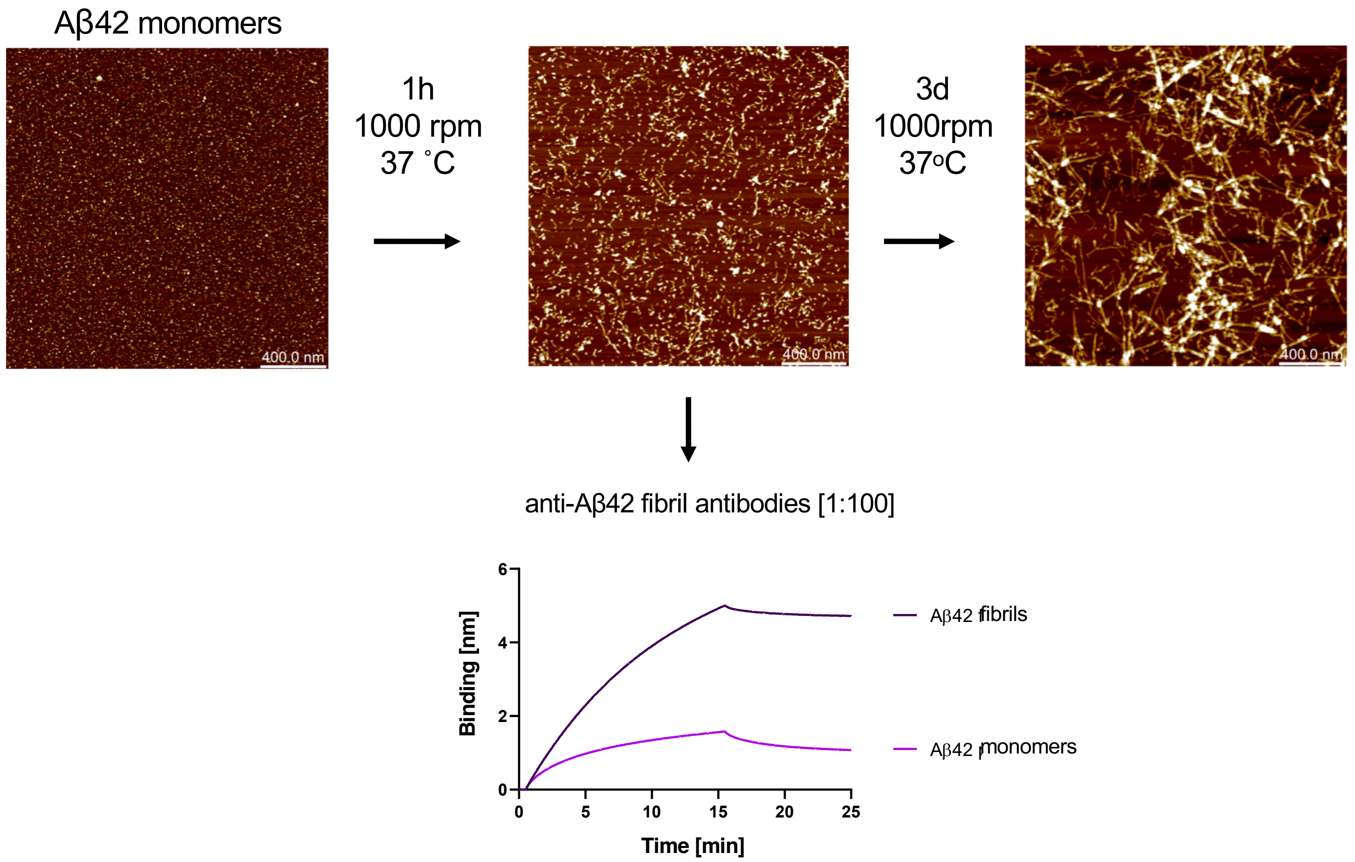


**Figure S23.** Formation of an Aβ42 micro-fibrils for a BLI biosensor immobilization. AFM images taken at three steps of the Aβ42 fibrils formation; (left) mixture of WT and biotinylated Aβ42 peptides (at 10:1 molar ratio) prior to fibril formation; (middle) micro-fibrils formed during 1 hour incubation with shaking at 37°C – these micro-fibrils were immobilized to the BLI sensor. (right) fibrils formed during 3 days of incubation with shaking at 37°C. (bottom) Immobilized Aβ42 fibrils, but not monomeric peptides, bind anti-amyloid antibodies.

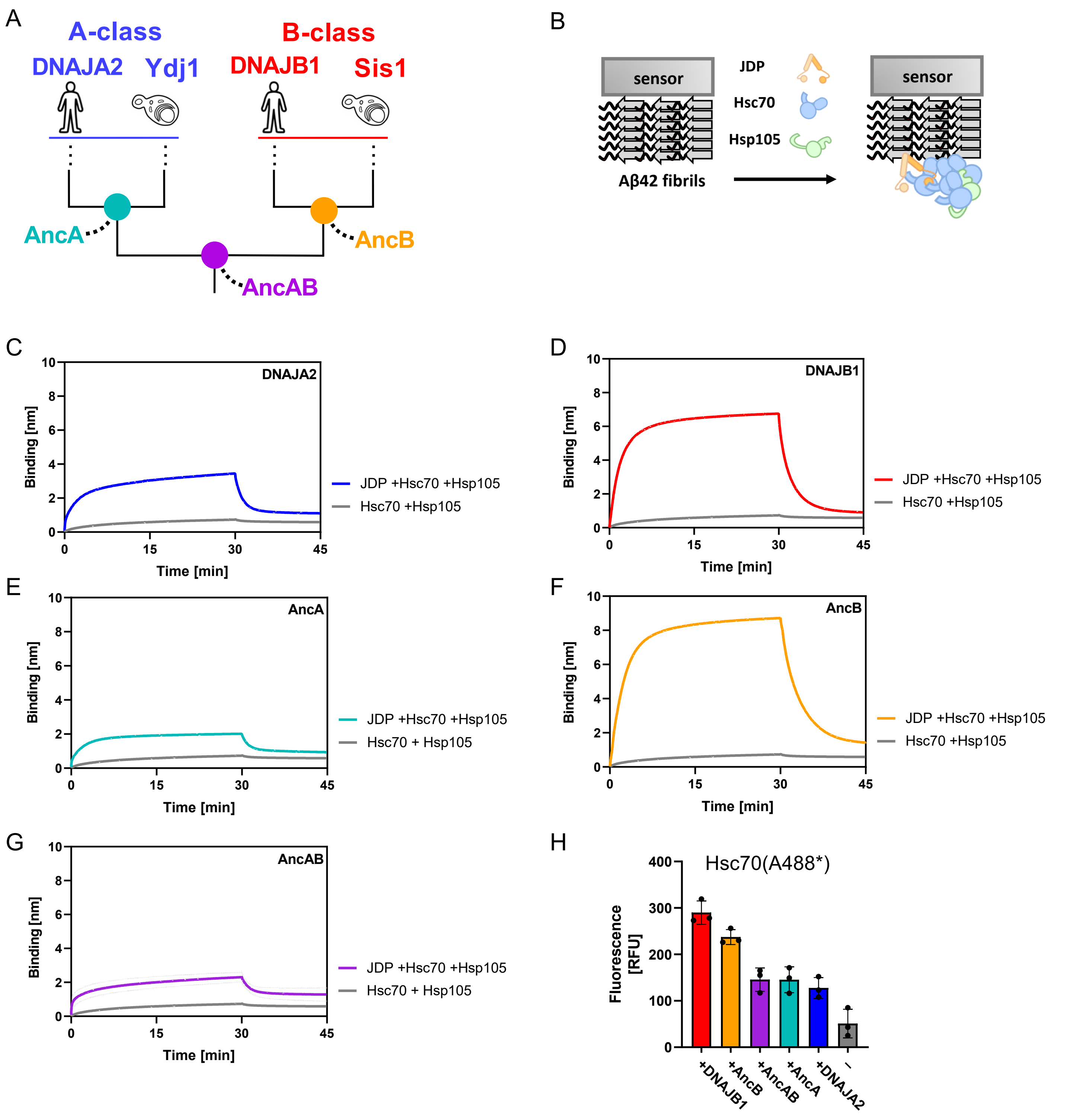


**Figure S24.** Ancestral JDPs' driven recruitment of Hsc70/Hsp105 to the sensor immobilized Aβ42 amyloid fibrils. (A) Simplified phylogeny of class A and $B^C$ JDPs. (B) The scheme of the Aβ42 binding experiment. (C-G) Interaction of Hsc70 1 µM and Hsp105 0.1 µM with BLI biosensor immobilized Aβ42 amyloid fibrils was monitored in the absence (grey) or presence (colored) of ancestral and contemporary JDPs 1 µM, as indicated. (H) Levels of Hsc70 interacting with Aβ42 amyloid fibrils were quantified using fluorescently labeled Hsc70 (A488*-Hsc70). Sensor with immobilized Aβ42 fibril was incubated for 30 minutes with a mixture of A488*-Hsc70 1 µM, Hsp105 0.1 µM and ancestral or contemporary JDP 1 µM, as indicated – association step. Fluorescence was measured after sensor was incubated for 15 minutes in a buffer without proteins -dissociation step.

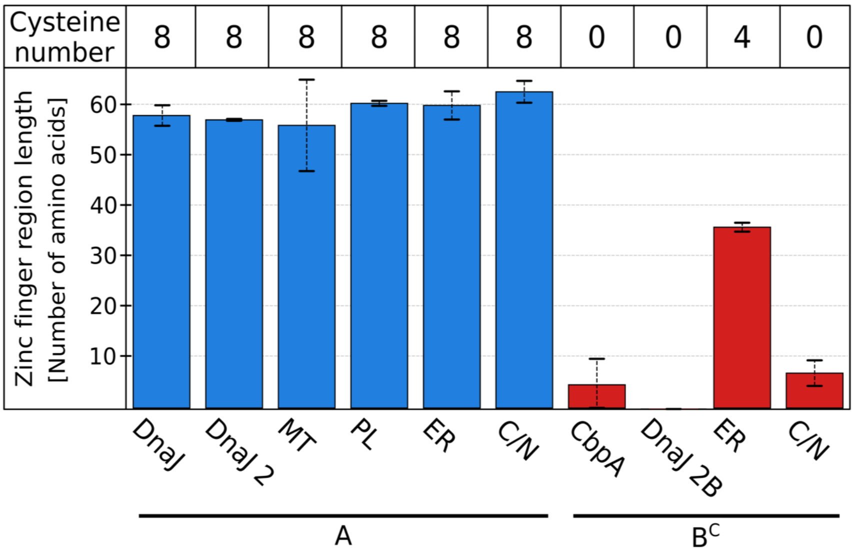


**Figure S25**. Distribution of the ZnF length and cysteine content across class A, $B^C$ and $B'^{(ST)}$ JDPs. The length of ZnF for class A (blue) and $B^C$ (red) JDPs. The number of cysteine residues within ZnF is indicated above the bars.
